# NF1 modulates microtubule repair and sensitivity to antibody-drug conjugates

**DOI:** 10.1101/2023.12.06.569572

**Authors:** Bruno Achutti Duso, Eleonora Messuti, Giulia Tini, Emanuele Bonetti, Alessia Castiglioni, Gianmaria Frigè, Giuseppe Ciossani, Silvia Monzani, Chiara Soriani, Daria Khuntsariya, Nicolò Roda, Andrea Polazzi, Marica R. Ippolito, Elena G. Doronzoro, Eltjona Mane, Martina Milani, Michele Bottosso, Martina Pagliuca, Alessia Farfalla, Ambra Dondi, Eleonora Nicolo’, Carolina Reduzzi, Costantino Jemos, Elena Guerini-Rocco, Simona Rodighiero, Daniela Tosoni, Arielle Medford, Andrew Davis, Paolo D’amico, Stefano Santaguida, Massimo Cristofanill, Marcus Braun, Zdenek Lansky, Luigi Scietti, Pier Giuseppe Pelicci, Luca Mazzarella

## Abstract

Antibody-Drug Conjugates (ADC) have revolutionized the treatment of several tumors, and extensive research is being devoted to the identification of predictive biomarkers. These are particularly sought after in fields, like breast cancer, in which multiple ADCs with identical target but different payloads have been approved. NF1 is a tumor suppressor widely mutated across several cancers, best characterized as an inhibitor of RAS signaling. Additional functions have been proposed but not deeply investigated, due to its large size and complex domain structure. Whether somatic NF1 mutations can be used to guide clinical decisions is not known.

Here, combining patient data, *in vitro*/*in vivo* models and protein biochemistry, we show that NF1 loss sensitizes cancer cells to T-DM1, the first approved ADC in breast cancer, through a novel, RAS-independent function on microtubular dynamics and repair. NF1 exhibits all biochemical properties of a bona fide Microtubule-Associated Protein (MAP) and specifically enhances intratubular repair, a recently discovered phenomenon whose regulation remains poorly characterized. NF1 loss results in mitotic defects and low-grade aneuploidy in cell lines and patients. Increased sensitivity to T-DM1 upon NF1 loss is confirmed in breast cancer patients analysed across institutions in Europe and USA.

Our results define NF1 as a key regulatory factor for microtubular repair and the first ADC payload-associated predictive biomarker identified to date.

## Introduction

Antibody-Drug Conjugates (ADC) have revolutionized the therapeutic landscape in oncology, leading to unprecedented survival advantages across multiple cancer types ^1,2^. In some tumors like breast cancer (BC), multiple ADCs have been approved and several more are in development, generating an increasing demand for biomarkers predictive of response to inform clinical decisions. Although all three components of ADCs (antibody, linker and payload) may be differently impacted by tumor/host biomarkers, research has so far focused mostly on antibody targets. However, there is increasing recognition that ADC activity is not simply determined by the levels of expression of their cognate antigen ^2^. No clear payload-associated predictive biomarker has been identified to date. In HER2+ BC, an area in which ADC development has been particularly disruptive, there are now two approved agents (Trastuzumab Emtansine, T-DM1, and Trastuzumab Deruxtecan, T-Dxd) and several in development, all targeting the same receptor (HER2), in some cases with the same antibody (Trastuzumab), but differing by linker and payload ^3^. Thus, rational treatment choices between ADCs are likely to be dictated by tumor-intrinsic differences in payload sensitivity.

Among payloads, derivatives of maytansine like DM1 have played a critical role, leading to the first ADC approval in solid tumors ^4^, and remain actively researched drugs. Maytansine was identified from the plant Maytenus Ovatus in the 70s but initially failed in clinical trials due to its low therapeutic index ^5^. Uniquely among microtubule-targeting agents, the maytansin binding site becomes inaccessible once tubulin is incorporated in the microtubule lattice, as it is positioned at the interface between two tubulin dimers ^6^. Thus, maytansine is currently thought to act by inhibiting de novo tubulin incorporation at the microtubule end ^7^. However, this model fails to explain why maytansinoids are vastly more effective in cells (in the nanomolar range) vs *in vitro* (in the micromolar range), suggesting that the cellular context includes factors able to enhance maytansine activity ^8^. Precise measurement of binding stoichiometry revealed a number of binding sites for maytansine in excess to the theoretical 13 exposed sites at each microtubule ^7^, but the nature of these additional binding sites is unknown.

Neurofibromatosis 1 (*NF1*) is among the first tumor suppressor genes identified ^9^. Loss-of-function (LoF) mutations in *NF1* are at the basis of the onco-developmental syndrome neurofibromatosis and are frequently found at the somatic level across multiple tumors ^10^. In BC, *NF1* LoF has been implicated in resistance to both endocrine and anti-HER2 agents. This has been attributed to the best characterized activity of NF1, its GTPase-activating protein (GAP) function that attenuates RAS signalling, or to a novel transcriptional regulatory activity ^11–13^. *NF1* encodes Neurofibromin (NF1), a large multifunctional protein consisting of 2,818 amino acids ^14^. Because of its large size and difficulty in cloning, NF1 structure has resisted characterization for several years, and only recently high-resolution reconstructions by Cryo-EM have been obtained ^15,16^. In early studies, NF1 was found to co-purify with tubulin in insect cells ^17^ and to co-localize with microtubules by immunofluorescence (IF) ^18^; two regions in NF1 have been proposed to mediate this interaction^19^. The physiological relevance of this interaction remains uncertain, and NF1 is not generally included among bona fide Microtubule-Associated Protein (MAP) ^20^.

Our understanding of microtubular dynamics has been significantly modified in recent years by the finding that dynamic changes do not only occur at the microtubule end (“dynamic instability”) but also within the lattice, where the mechanical stress induced by motors or other sources generates discontinuities that are continuously repaired by intra-lattice tubulin incorporation ^21,22^. To date, only two MAPs, CLASP and SSNA1 ^23,24^ have been found to recognize and modulate such microtubule damage. None of these proteins is commonly mutated in cancer, so the relevance of microtubule repair for tumorigenesis remains uncertain, despite the established relevance of microtubule-dependent functions such as mitosis.

Here, we show that NF1 loss is a common event in metastatic HER2+ BC and is associated with resistance to several agents but increased sensitivity to the ADC T-DM1. This sensitization is specifically due to a RAS-independent activity of NF1 on the modulation of microtubule damage repair.

## Results

### 1. *NF1* is the most differentially mutated gene in metastatic *vs* primary HER2+ BC and its loss leads to differential sensitivity to HER2-targeting agents

To identify genes potentially involved in resistance to conventional anti-HER2 agents in breast cancer, we compared the frequency of mutation occurrence between metastatic and primary HER2-amplified BC, since virtually all metachronous metastases develop after exposure to targeted agents, typically at least Trastuzumab. We pooled publicly available datasets of BC sequencing TCGA and AACR-GENIE ^25,26^ and identified primitive (n=1066) *vs* metastatic (n=1418) cases. We compared frequencies of mutated genes in the overall cohort and in the *ERBB2*-amplified subgroup, identified based on copy number data rather than histological HER2 status, which is often unreported or unreliable. *NF1* was identified as the most significant differentially mutated gene between metastatic and primary HER2+ BC (OR=5.14, pvalue=2.93 x 10^-4^, adj pvalue=0.01, figure 1A); *NF1* was also more frequently mutated in metastatic than in primary in the overall cohort, but this was comparatively less significant (OR=1.739, pvalue=0.00249, adj pvalue=1, supplementary figure 1A). The mutational spectrum in metastatic tumors was dominated by nonsense mutations and missense mutations scattered throughout the gene, indicative of loss of function and typical of a tumour-suppressor gene (figure 1B and supplementary figure 1B).

**Figure 1.**
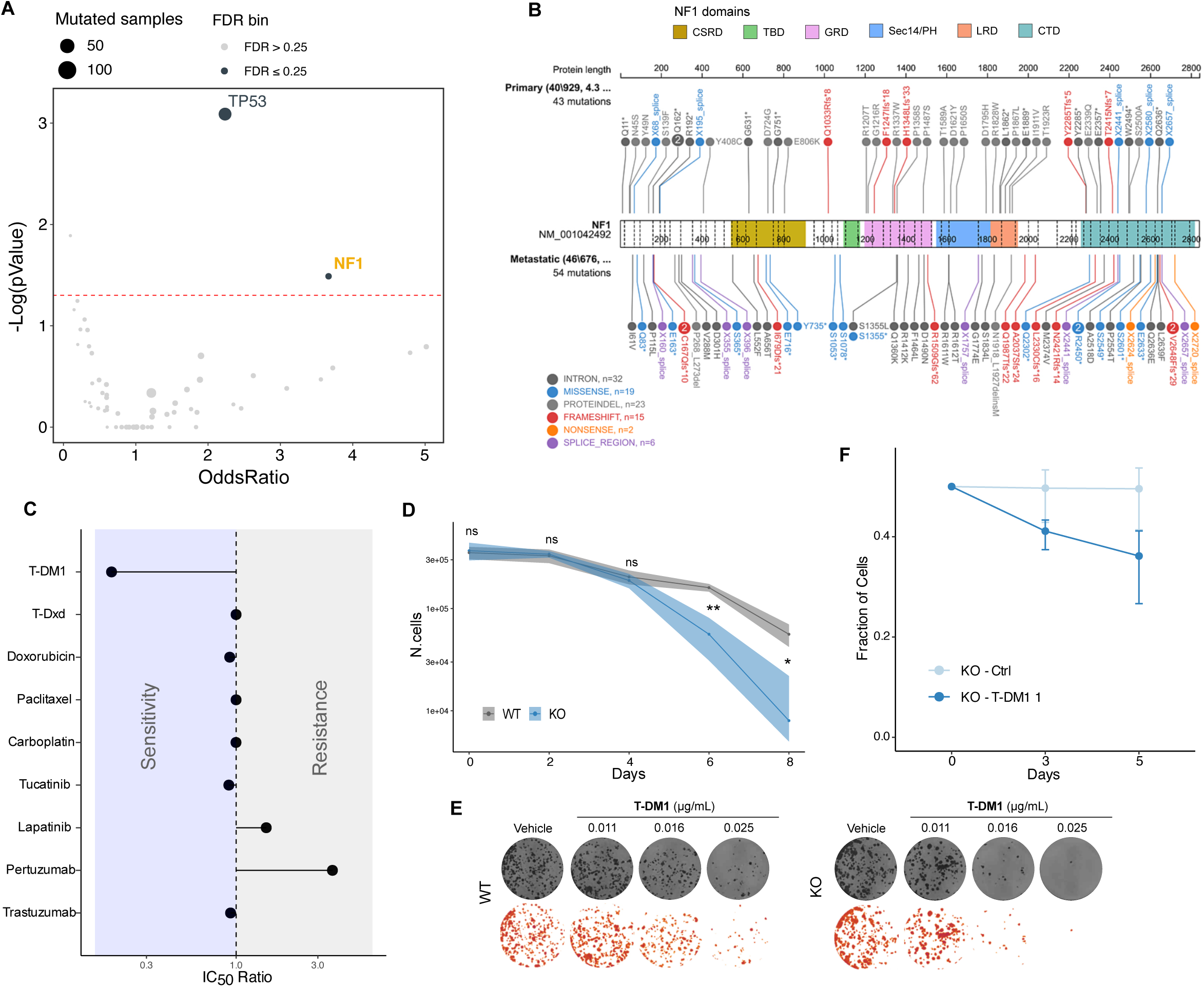
NF1 is differentially mutated in metastatic vs primary HER2+ BC and its loss leads to differential sensitivity to anti-HER2 agents. **A)** Comparison of mutation frequency in unmatched metastatic vs primary tumors in ERBB2-amplified BC patients, collected from TCGA and AACR-GENIE. **B)** Mapping of point mutations in metastatic vs primary breast tumors in ERBB2-amplified patients. **C)** Ratio of IC50 values (log10 scale) in short-term (3 days) BrdU incorporation assays between BT-474^KO^ and BT-474^WT^ cells exposed to the indicated agents. **D)** 8-day growth curves BT-474^KO^ and BT-474^WT^ cells treated with T-DM1 1 μg/ml. To draw the ribbon in log10 scale, non-positive lower boundaries (time-point 8, KO) were substituted by the value 5000. Post-hoc pairwise t-test (after significant 2-way ANOVA for time series (see methods)) are reported (ns p≥0.05; *p<0.05, **p<0.001) **E)** Long-term (14 days) colony formation assay for BT-474^KO^ and BT-474^WT^ cell lines treated with T-DM1 at the indicated doses. **F)** Co-culture of H2B-GFP-BT-474^WT^ or H2B-mCherry-BT-474^KO^ cells; mean ± SD; paired 2-tailed t-test. * p < 0.05, ** p < 0.01

To investigate the biological consequences of *NF1* loss in HER2+ BC, we modelled NF1 loss of function by CRISPR-Cas9 mediated generation of *NF1* knockout (KO) clones of the widely used HER2+ BC cell lines BT-474, SK-BR-3 and HCC1954 (supplementary figure 1C-D). We then subjected BT-474^WT^ and BT-474^KO^ cells to a pharmacological screen with all drugs currently approved for HER2+ BC treatment, including chemotherapeutics, monoclonal antibodies, Tyrosine-Kinase Inhibitors (TKI) and antibody-drug conjugates (ADC). We decided to use a 4-day BrdU incorporation assay, which we found to have a wider dynamic range for short-term assays with slow-proliferating cells like BT-474. For many agents, in particular pertuzumab, IC_50_ values were higher, supporting the idea that *NF1* loss is associated resistance to standard HER2-targeting agents, in agreement with results of a recent study in which NF1 levels were lowered by RNA interference ^12^ (figure 1C and supplementary figure 1E). Unexpectedly, NF1 loss was not only associated with resistance: the ADC T-DM1 was more effective in BT-474^KO^ cells, which exhibited a substantially lower IC50. This differential sensitivity became more apparent in longer term growth curves (figure 1D), colony formation assays (figure 1E) and in coculture experiments in which BT-474^WT^ and BT-474^KO^ cells were differentially labelled with mCherry and GFP respectively (figure 1E and supplementary figure 1F). Differential sensitivity was confirmed in HCC1954 cells (supplementary figure 1G-I).

### 2. NF1 loss leads to increased efficacy of T-DM1 treatment in vitro, in vivo and in patients

In BT-474^KO^ cells, T-DM1 led to enhanced G2/M arrest compared to BT-474^WT^ cells (figure 2A-C and supplementary figure 2A), and induction of apoptosis markers (cleaved PARP1, figure 2D). Furthermore, residual cells after T-DM1 treatment stained more strongly with the senescence marker beta-galactosidase in BT-474^KO^ vs BT-474^WT^ cells (supplementary figure 2B-C).

**Figure 2.**
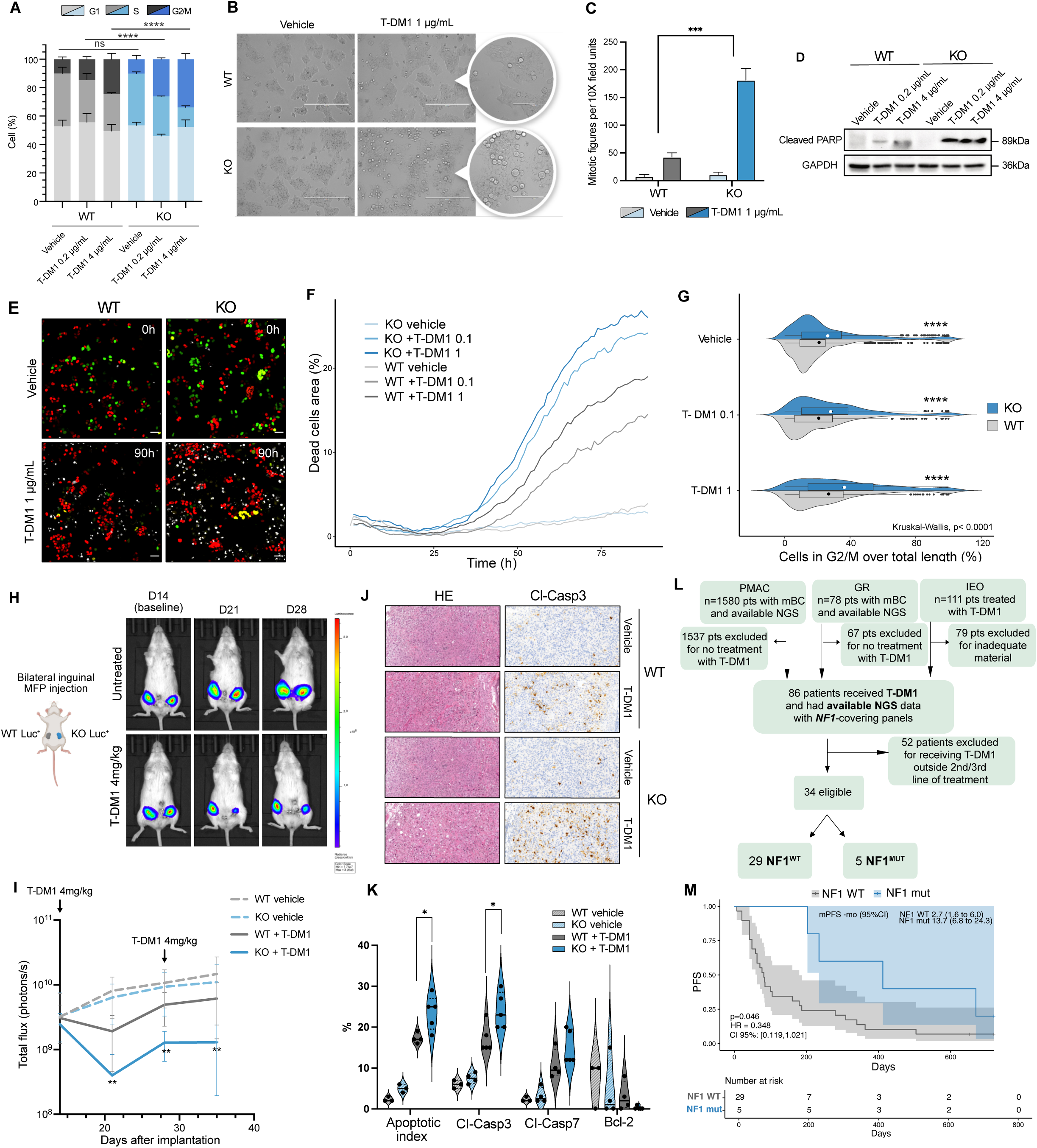
NF1 loss leads to increased G2/M arrest and apoptosis upon T-DM1 treatment. **A)** Propidium-iodide (PI) cell cycle analysis in BT-474^WT^ and BT-474^KO^ cells after 36h of T-DM1 at the indicated doses; **** p < 0.0001, χ2 test; error bars are SD. **B)** Round-up figures on BT-474^KO^ and BT-474^WT^ cells, indicative of ongoing terminal mitosis, observed in phase contrast microscopy after 24h of T-DM1 treatment. **C)** Quantification of mitotic figures per field of view, *** p < 0.001, Fisher’s exact test; error bars are SD. **D)** Western blot of BT-474^KO^ and BT-474^WT^ cells for cleaved PARP at 48h after T-DM1 treatment. **E)** Cell cycle analysis by FUCCICa-transduction of BT-474^KO^ and BT-474^WT^ cells. Cells transition through red (G1), green (S), yellow (G2/M) and white (DRTM - a live-cell impermeable, far-red emitting DNA dye that marks for cell death). Cells were live-imaged for 90 hours every 10 minutes after addition of T-DM1 1 mg/ml. Representative frames 0h and 90h are shown. **F)** Cell viability (% area covered by DRTM signal) over time of BT-474^KO^ and BT-474^WT^ cells treated with T-DM1. **G)** Distribution of fraction of time spent in G2/M upon T-DM1 treatment. Kruskal-Wallis with Dunn’s post-hoc analysis between WT and KO. **** p value<0.0001. **H)** Bioluminescent images and **I)** Growth of HCC1954^WT^ and HCC1954^KO^ xenograft mammary tumors treated biweekly with vehicle or T-DM1 (4 mg/kg); n = 10 mice per group, means of total flux (photons/s) with SD are plotted. **J)** Hematoxylin-eosin (HE) and Cleaved Caspase 3 staining of tumor masses isolated 24 hours after T-DM1 treatment. **K)** Quantification of data in J. One-way ANOVA with Tukey post-hoc analysis. *p<0.05. **L)** Flow diagram of patient selection strategy and NF1 status assessment. **M)** Progression Free Survival (PFS) analysis for patients. Log-rank p=0.046. Hazard Ratio (HR), 95% confidence intervals (CI), risk table and median PFS (mPFS) are reported.

To further characterize cell cycle dynamics, we engineered BT-474^WT^ and BT-474^KO^ to express the FUCCI(Ca) reporter ^27^, which allows the monitoring of G1, S and G2M phases in live cell imaging experiments; the DRAQ7 viability dye was added to monitor cell death. The kinetics of cell death was similar in BT-474^WT^ and BT-474^KO^ cells, initiating ∼ 30 hrs after T-DM1 administration (figure 2E-F), with death typically occurring upon exit from M phase or right after re-entry in G1 (supplementary figure 2D-E and supplementary video 1-2. T-DM1-treated BT-474^KO^ cell exhibited significantly prolonged G2/M phase compared to BT-474^WT^ cells (figure 2G). Then, we tested if such differential activity could also be observed *in vivo.* We chose the HCC-1954 model as it provides faster, more reliable and estrogen-independent growth kinetics compared to BT-474 and SK-BR-3. We injected luciferase-expressing HCC-1954^WT^ and HCC-1954^KO^ cells in the two opposite flanks of Nod-Scid-IL2GR^KO^ (NSG) mice, which we randomized to treatment by either saline or T-DM1 at days 15 and 29 after tumor injection. Vehicle-treated tumors did not exhibit differential growth, whereas T-DM1 resulted in significantly stronger growth inhibition of HCC-1954^KO^ vs HCC-1954^WT^ tumors (figure 2H-I). Histological analysis of tumors retrieved 48 hours after T-DM1 treatment revealed a significantly higher apoptotic index and cleaved Caspase 3 in HCC-1954^KO^ masses (figure 2J-K).

Finally, we assessed the interaction between NF1 status and progression-free survival (PFS) in 34 HER2+ BC patients who received T-DM1 as second- or third-line therapy in the metastatic setting, identified retrospectively across 5 centers in Europe and USA. The patients’ main characteristics are reported in supplementary table 1; NF1 mutations in supplementary table 2. Mean PFS was 13.7 months (95% CI, 6.8 to 24.3) for the NF1^MUT^ group and 2.7 months (95% CI, 1.6 to 6) for the NF1^WT^ group (P = 0.04) with a hazard ratio of 0.37 (95% CI, 0.17 to 0.81, figure 2M-N).

We conclude that genetic ablation of NF1 leads to increased sensitivity to T-DM1 *in vitro*, *in vivo* and in patients

### 3. NF1 loss-associated hypersensitivity to T-DM1 is due to the DM1 payload and is not reproduced by RAS hyperactivation

We then investigated if differential response to T-DM1 is due to biological factors affecting antibody binding/internalization or response to the DM1 payload. First, BT-474^KO^ cells did not exhibit higher levels of HER2 expression (supplementary figure 3A). Also, IF for endocytosed T-DM1 did not reveal any major difference in ADC internalization (supplementary figure 3B-C). Conversely, treatment with the cell-permeable naked payload DM1 was more effective in in BT-474^KO^ compared to BT-474^WT^ cells in long-term growth curves and colony-formation assays (figure 3A-B), similarly to T-DM1 and differently from Trastuzumab alone. Upon treatment with T-Dxd, the other approved ADC in HER2+ BC, no difference could be observed between WT and KO cells by internalization nor drug response in long-term cultures, co-cultures or colony formation (supplementary figure 3B-F). Similar results were obtained on HCC1954 cells (supplementary figure 3 G). DM1 was also the only microtubule-targeting agent, among the many tested (vinblastine, paclitaxel, MMAE, Paclitaxel, Pironetin, supplementary figure 3H), exhibiting selectivity for NF1-KO cells.

**Figure 3.**
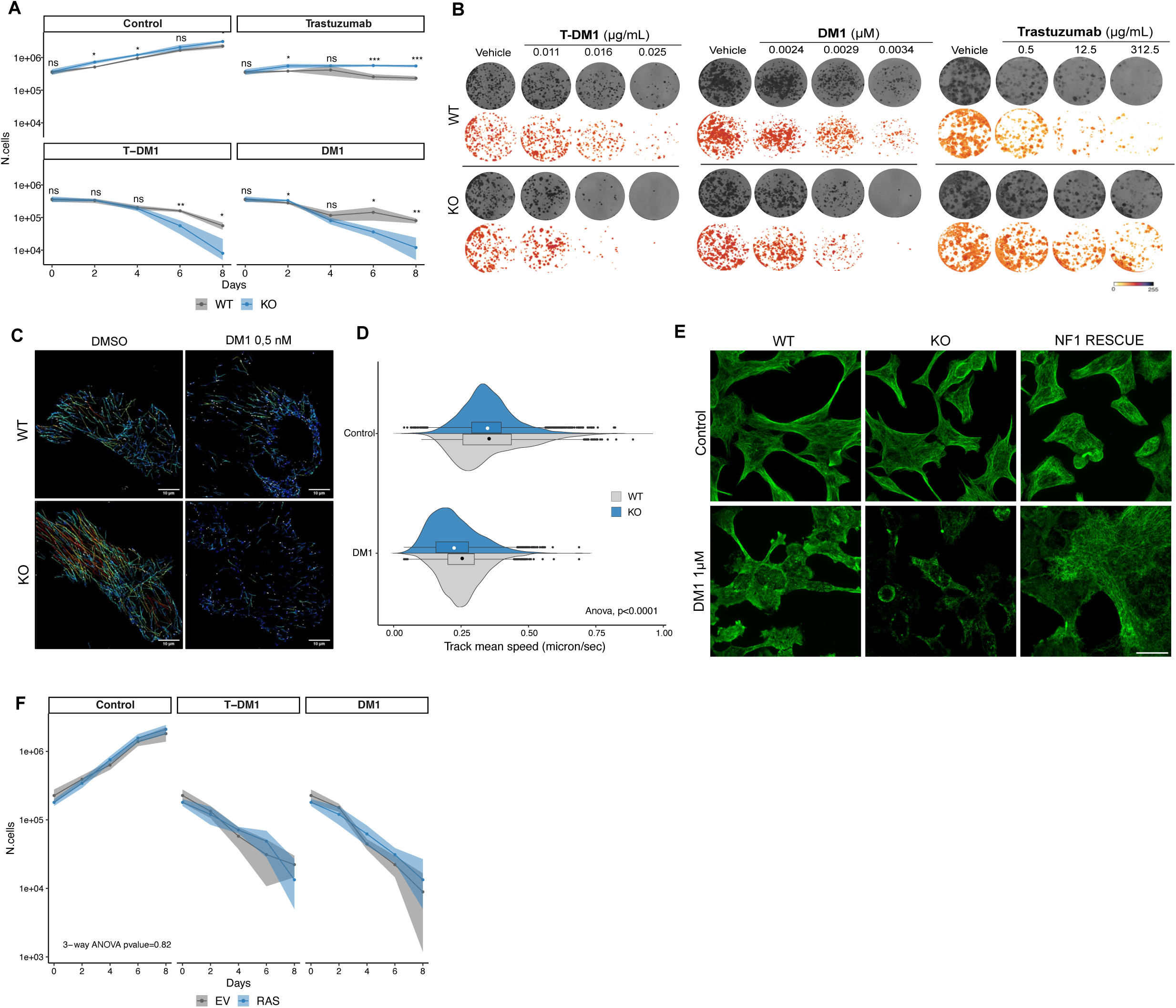
NF1 loss-associated hypersensitivity to T-DM1 is due to the DM1 payload and is RAS-independent. **A)** 8-days growth curves on BT-474^WT^ and BT-474^KO^ cells treated with DM1 25nM, T-DM1 1 μg/ml and Trastuzumab 100 μg/ml. Rituximab, a human IgG1 monoclonal antibody, and DMSO were used as control. To draw the ribbon in log10 scale, non-positive lower boundaries (time-point 8, KO, DM1 and TDM1) were substituted by the value 5000. Post-hoc pairwise t-test (after significant 3-way ANOVA for time series (see methods)) are reported (ns p≥0.05; *p<0.05, **p<0.001, ***p<0.0001). **B)** Colony formation assay for BT-474^WT^ and BT-474^KO^ cell lines treated with T-DM1, DM1 and Trastuzumab at the indicated doses. **C)** Reconstruction of microtubule tip trajectories after monitoring of microtubule elongation in HCC1954^WT^ and HCC1954^KO^ cells transiently transfected with an EB3-GFP-expressing vector and monitored by live cell imaging for 1 minute with acquisitions every 4 seconds. Microtubule tip fates were reconstructed by the TrackMate ImageJ plugin (scale bar= 10 μm). **D)** Quantification of tip mean speed. At least 1000 events from at least 3 cells were calculated per condition. 2-way ANOVA with Tukey’s HSD. **** p < 0.001. **E)** Immunofluorescence of HCC1954^WT^ and HCC1954^KO^ cells treated with DM1 1 μM for 45 minutes stained for tubulin (green). Scale bar= 30 μm. **F)** 8-days growth curves with DM1 25nM, T-DM1 1 μg/ml on BT-474^EV^ and BT-474^KRAS^ ^G12V^ cells. To draw the ribbon in log10 scale, non-positive lower boundaries (time 8, RAS OE, DM1 and TDM1) were substituted by the value 5000. 3-way ANOVA tests for time series not significant.

We analyzed in detail the kinetics of microtubule depolymerization in response to DM1 and how this is affected by NF1. We engineered BT474^WT^ and BT474^KO^ cells to express the EB3-GFP reporter, which binds to microtubular plus-ends and allows their tracking by live cell imaging and the computation of MT dynamics parameters in living cells ^28^. In the absence of treatment, average microtubule growth speed was not significantly different between BT-474^KO^ and BT-474^WT^ cells, but upon DM1 treatment, microtubule speed was significantly reduced in BT-474^KO^ cells (figure 3C-D and supplementary video 3-6). To further confirm the role of NF1 in modulating DM1-dependent microtubule depolymerization, we analyzed HCC-1954 cells, in which we managed to obtain stable re-expression of WT NF1 in a KO background by lentiviral transduction (supplementary figure 3I). NF1 ablation or rescue did not lead to visible differences in microtubule signal in the absence of treatment. Upon addition of DM1, microtubules were visibly more depolymerized in HCC- 1954^KO^ cells; this was completely rescued upon NF1 re-expression (figure 3E).

We then investigated if hypersensitivity to T-DM1 is reproduced by oncogenic RAS signaling activation, which is often phenocopied by NF1 loss. We generated BT-474 cells expressing oncogenic p.G12V KRAS (BT-474 ^G12V^). These cells exhibited the expected RAS hyperactivation, as revealed by RAS-GTP pulldown assays, similar or higher than that achieved by NF1 loss (supplementary figure 3J). We did not observe any major difference in long-term growth either in unperturbed conditions or upon treatment with T-DM1 or DM1 (figure 3F).

We conclude that increased sensitivity to T-DM1 upon NF1 loss is due specifically to the DM1 payload and is not a consequence of differential target-mediated drug bioavailability or RAS hyperactivation.

### 4. NF1 loss leads to mitotic aberrations and aneuploidy

To further characterize the effects of *NF1* loss and T-DM1 treatment, we analyzed the transcriptome of BT-474^KO^ and BT-474^WT^ cells subjected to T-DM1 treatment. Principal component analysis (PCA) of RNAseq showed clear segregation of biological replicates with genotype and treatment (figure 4A). When we compared transcriptional changes induced by T-DM1, we identified a set of 256 genes selectively or more markedly downregulated in BT-474^KO^ compared to BT-474^WT^ cells (figure 4B). Gene Ontology analysis on this gene set revealed enrichment for terms related to mitosis and mitotic spindle (figure 4C and supplementary table 2), further suggesting a role for NF1 in modulating microtubules and mitosis.

**Figure 4.**
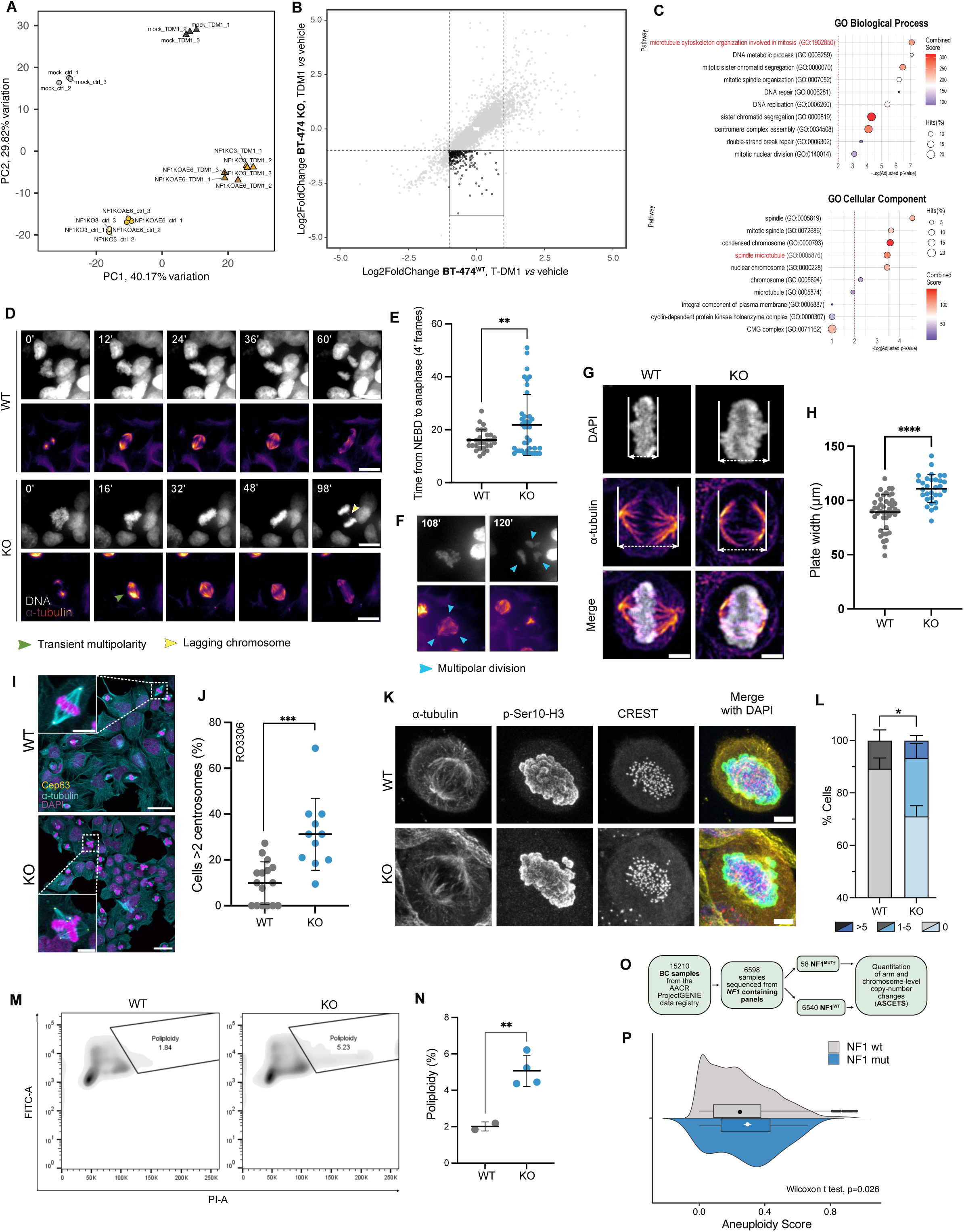
NF1loss leads to mitotic aberrations and aneuploidy. **A)** Principal Component Analysis of RNAseq of BT-474^WT^ and BT-474^KO^ cells treated with T-DM1 at 1 μg/ml for 36 hours (3 independent biological replicates per condition). Two independently derived KO clones were analysed. **B)** Comparison of transcriptional changes from RNAseq analysis, expressed as log2 fold change of treated vs untreated, per genotype. Highlighted genes are significantly more downregulated in BT-474^KO^ in response to T-DM1. **C)** Gene ontology analysis of genes highlighted in panel B. Spindle/microtubule-associated pathways are highlighted in red. **D)** Representative frames of live cell imaging of BT-474^WT^ and BT-474^KO^ cells stably expressing H2B-GFP or H2B-mCherry respectively and monitored over 100 minutes, with acquisitions every 4 minutes (scale bar=15 μm). Cells were incubated with cell-permeable, fluorescent Sir-tubulin to monitor spindle formation. Arrows indicate lagging chromosomes. DAPI in grey and tubulin in red. **E)** Quantification of mitosis length, indicated as time from Nuclear Envelope BreakDown (NEBD) to anaphase end (mean ± SD are represented, Wilcoxon test, **p<0.01). **F)** Representative frames of a BT-474^KO^ cell engaging anaphase in a multipolar conformation. DAPI in grey and tubulin in red. **G)** Representative images of mitotic plate measurement in BT-474^WT^ and BT-474^KO^ cells (scale bar=5 μm). **H)** Quantification of plate width (mean ± SD are represented, Wilcoxon test, ****p<0.0001). **I)** Representative images of HCC1954^WT^ and HCC1954^KO^ cells synchronized by RO3306. Cells are stained with anti-alpha tubulin, centrosome marker Cep63 and DAPI. **J)** Fraction of cells with supernumerary centrosomes per field of view (mean ± SD are represented, Wilcoxon test, ***p<0.001). **K)** Representative images of chromosome missegregation (arrowhead) in BT-474^WT^ and BT-474^KO^ metaphase synchronized cells (scale bar=5 μm). The centromere marker CREST is used to identify centromeres. **L)** Quantification of data from experiment in K (mean ± SD are represented, Wilcoxon test, *p<0.05). **M)** Representative FACS plots of cell cycle analysis by propidium iodide and BrdU staining in BT-474^WT^ and BT-474^KO^ cells. Polyploid cells are gated. **N)** Quantification of data in panel M (mean ± SD are represented, Wilcoxon test, **p<0.01). **O)** CONSORT diagram of breast cancer patient selection strategy for aneuploidy score analysis from AACR GENIE dataset. NF1-mutated patients were selected from the AACR Genie cohort for the presence of variants pathogenic according to alpha missense. **P)** Quantification of the ASCETS-computed aneuploidy score in NF1-mutated vs -nonmutated patients. *p<0.05 by Wilcoxon test.

To investigate in detail this hypothesis, we monitored mitosis duration and anomalies using live cell imaging of BT-474^WT^ and BT-474^KO^ cells engineered to express H2B-GFP and H2B-mCherry, respectively, to follow mitotic chromatin condensation, and exposed to fluorescent SiR-tubulin to monitor spindle formation. BT-474^KO-H2BmCherry^ cells exhibited significantly prolonged mitosis, lagging chromosomes and pseudodipolar mitoses (figure 4D-F and supplementary video 7-8) compared to BT-474^WT-H2BGFP^. BT-474^KO^ cells showed an enlarged mitotic plate (figure 4G-H), suggestive of chromosomal instability (CIN) ^29^. Immunofluorescence for tubulin and centrosome marker CEP63 cells synchronized by the CDK1 inhibitor RO3306 exhibit a significantly higher prevalence of cells with supranumerary centrosomes (figure 4I-J and supplementary), which lead to pseudo-dipolar mitoses in HCC1954^KO^ over HCC1954^WT^ cells. These results were confirmed in SKBR3 cells synchronized by double thymidine block (supplementary figure 4A-B). Since altered mitosis is associated with the acquisition of chromosomal aberrations, we measured the impact of NF1 loss on aneuploidy. Upon synchronizations, HCC1954^KO^ cells exhibited an increased rate of chromosome misalignment (figure 4K). BT-474^KO^ had a higher fraction of cells exhibiting >2n DNA content in cell culture as measured by cell cycle analysis (figure 4M-N). To verify if NF1 loss is a clinically relevant determinant of tumor aneuploidy, we computed the aneuploidy score (AS) ^30^ for tumor samples from breast cancer patients in the AACR Genie cohort ^25^. As shown in figure 4O, patients with an NF1 mutation predicted to have a pathogenic impact according to Alpha Missense ^31^ showed a significantly higher AS compared to those without.

We conclude that NF1 loss leads to important mitotic aberrations that favour the acquisition of aneuploidy.

### 5. NF1 is a microtubule-associated protein (MAP)

The impact of NF1 loss on mitosis suggested a direct role for NF1 on microtubule dynamics. To study the biochemistry of NF1 interaction with tubulin and microtubules, we produced recombinant NF1 in insect cells (supplementary figure 5A-B). As previous studies showed that the histidine tag used for purification may generate spurious interactions with microtubules^32^, we modified a previously generated NF1 expression vector^33^ by introducing a PreScission cleavage site (supplementary figure 5C). Purified full-length NF1 exhibited the expected biochemical properties as i) it formed dimers of ∼654 kDa as analyzed by mass photometry (supplementary figure 5D) and ii) it exhibited GTPase-Activating Protein (GAP) activity on RAS (supplementary figure 5E). To test whether NF1 directly binds to microtubules, we performed a classical microtubule co-sedimentation assay. Purified NF1 was fully soluble, but incubation with polymerized tubulin led to NF1 co-sedimentation with microtubules in the insoluble pellet (figure 5A). In turbidity-based tubulin polymerization assays, purified NF1 led to a significant acceleration of the polymerization curve, with a loss of the initial nucleation phase (figure 5B). NF1 decreased the critical tubulin concentration, i.e., the minimal concentration to initiate polymerization (supplementary figure 5F). Microscopic inspection of microtubules formed in the presence of NF1 revealed the formation of microtubule bundles (figure 5C), a property shared by several MAPs ^20^ which may also contribute to turbidity increase and complicate the interpretation of bulk polymerization assays. To remove the effect of bundle formation, we assessed the dynamics of individual microtubules by Total Internal Reflection Fluorescence (TIRF) microscopy on microtubules polymerizing from glass-immobilized stabilized microtubule seeds. The addition of NF1 led to a dose-dependent increase in the fraction of polymerizing microtubules, elongation speed and rescue frequency (figure 5D-G and supplementary videos 9-11).

**Figure 5.**
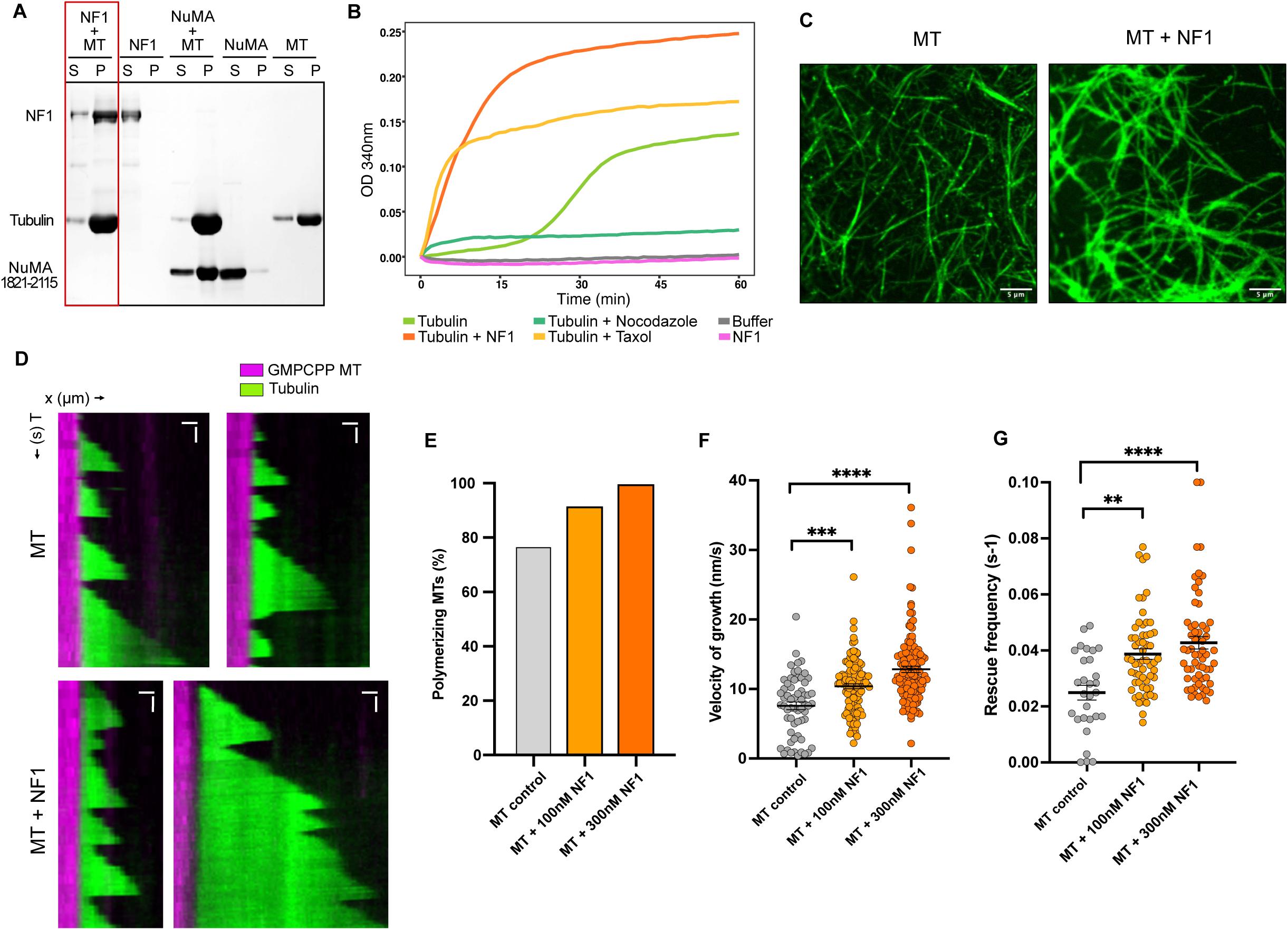
NF1 is a microtubule-associated protein (MAP) **A)** Co-sedimentation of NF1 with Taxol-stabilized microtubules. NuMa, a known microtubule-binding protein, was used as positive control. S=supernatant, P=pellet. Representative of 3 independent experiments. **B)** Turbidity-based tubulin polymerization assay. Tubulin (20 μM) was incubated alone or in combination with drugs or NF1 as indicated at 37°C. Polymerization was followed by continuous (1 reading per minute) reading of absorbance at 340nm over 60 minutes. **C)** Representative images of taxol-stabilized rhodamine-labeled microtubules incubated with or without NF1. **D)** Representative kymographs of microtubules grown with or without NF1 from GMPCPP-stabilized seeds and followed by TIRF microscopy. Scale bars: 2 μm (horizontal) and 60 s (vertical) **E)** Percentage of polymerizing microtubules, calculated as the number of GMPCPP seeds from which microtubules polymerized divided by the total number of seeds present in the acquired ROI. **F)** Velocity of microtubule growth (mean ± SEM from one over three independent experiments are represented, Kruskal Wallis test with Dunn post-hoc test, *** p<0.001, **** p<0.0001). **G)** Rescue frequency, calculated by dividing the number of rescues observed by the total shrinkage time (mean ± SEM from three independent experiments are represented, Kruskal Wallis test with Dunn post-hoc test, ** p<0.01, **** p<0.0001).

### 6. DM1 leads to intratubular damage that is repaired more efficiently in the presence of NF1

To characterize how NF1 loss modulates the pharmacodynamics of DM1 in cells, we performed Cellular Thermal Shift Assay (CETSA), which showed a significant right shift of the melting curve in BT-474^KO^ cells, indicating that DM1 engages tubulin more efficiently in BT-474^KO^ cells than in BT-474^WT^ (figure 6A).

**Figure 6.**
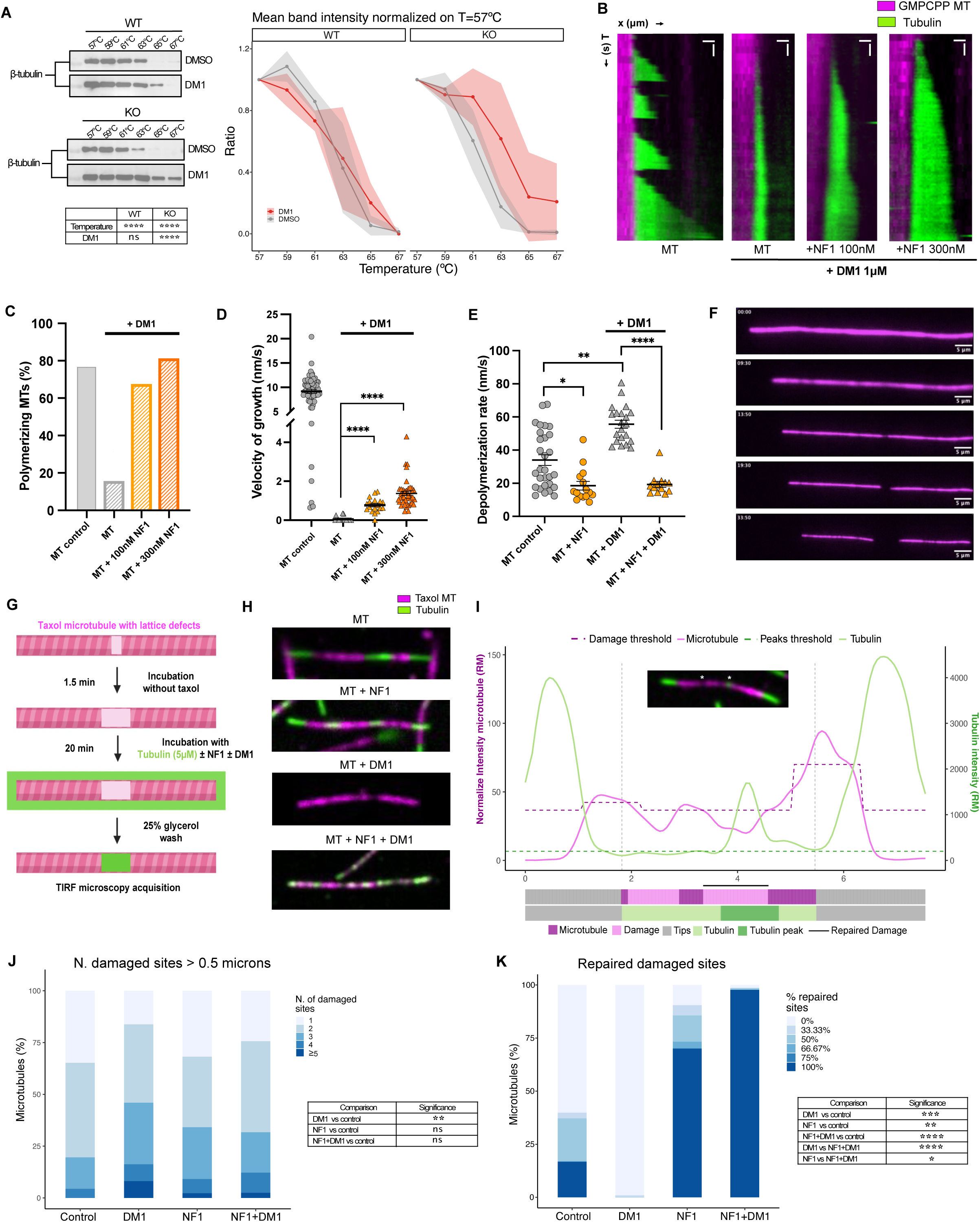
DM1 leads to intratubular damage that is repaired more efficiently in the presence of NF1. **A)** Western blot for β-tubulin of cell lysates from BT-474^WT^ and BT-474^KO^ cells, treated with 1 μM DM1 for 2 hours and then exposed to increasing temperatures as indicated (one over three independent experiments is represented). On the right, quantification of band intensities from CETSA western blots (n = 3 independent experiments), normalized on baseline signal at 57°C. Mean ± SD are indicated as dots and ribbons. 2-way ANOVA was used to assess significance of temperature and DM1 addition. **B)** Representative kymographs of microtubules grown with or without NF1 at indicated concentration and 1μM DM1 from GMPCPP-stabilized seeds and followed by TIRF microscopy. Scale bars: 2 μm (horizontal) and 60 s (vertical). **C)** Percentage of elongating microtubules, calculated as the number of GMPCPP seeds which polymerized microtubules divided by the total number of seeds present in the acquired ROI. **D)** Velocity of microtubule growth (mean ± SEM from three independent experiments are represented, Kruskal Wallis test with Dunn post-hoc test, **** p<0.0001). **E)** Depolymerization velocity of microtubules grown with or without 100nM NF1 and with or without 1μM DM1. Data are means ± SEM, Kruskal Wallis test with Dunn post-hoc test was used to assess significance **** p<0.0001. **F)** Single frames of time-lapse video of a taxol-stabilized rhodamine-labeled microtubule incubated with 1μM DM1 (scale bar=5 μm). **G)** Experimental design of microtubule repair experiment. **H)** Representative images showing Alexa 488-tubulin (green) incorporation sites into rhodamine-labeled microtubule lattice (magenta) with or without 300nM NF1 and with or without 1μM DM1. **I)** Representative plot of microtubule (magenta) and tubulin (green) signal intensity. Thresholds for the identification of damages and tubulin peaks are reported as dotted lines (violet and dark green). Horizontal bars below the intensity plot represent microtubule tips (grey), intact microtubule and damage sites (magenta); tubulin and its peaks (green). Damaged sites repaired by tubulin are highlighted with a black line. **J)** Distribution of damaged sites per microtubule within each experimental condition. Permutation test was used to assess significance. **K)** Percentage of repaired damaged sites for microtubules in each condition. Permutation test was used to assess significance.

We then investigated microtubules treated with DM1, with or without NF1, using our TIRF microscopy assay. DM1 led, as previously described ^7^, to a reduction in the fraction of elongating microtubules and in elongation speed (figure 6B-D). Addition of recombinant NF1 dose-dependently inhibited DM1 activity and allowed microtubular polymerization to occur, to the point that at 300 nM NF1 the fraction of polymerizing microtubules was completely rescued (figure 6B-C and supplementary videos 12-14). In conditions favoring depolymerization (low free tubulin concentration) of taxol-stabilized microtubules, DM1 accelerated depolymerization rate as expected. Addition of NF1 significantly reduced depolymerization rate both in control and treated microtubules (figure 6E). Additionally, we observed the appearance of areas of reduction in tubulin density in the microtubule lattice upon DM1 treatment, in some cases leading to microtubule breakage (figure 6F). Microtubule breakage was never observed in the presence of NF1. These findings suggest that DM1 is able to bind to intra-lattice sites, in addition to its known ability to bind dimeric tubulin or microtubule tips ^8^, causing or amplifying intratubular damage which may be modulated by NF1. We directly investigated lattice damage by adapting a method previously used in the analysis of the microtubule lattice repair ^23^. Upon formation of taxol-stabilized rhodamine-labelled microtubules, and a brief incubation in a buffer without taxol, which increases the size of possible lattice defects, free A488-labeled tubulin was added to the measurement chamber. A488-labeled tubulin incorporates in defects existing in the microtubule lattice and can be thus used to map sites of de novo tubulin incorporation (figure 6G-H). We developed a signal analysis method to identify areas of decreased tubulin intensity in the stabilized microtubule (“damaged site”), independently from the subsequent de novo incorporation (figure 6I and supplementary figure 6A). Our method allowed us to asses microtubule repair as the amount of *de novo* incorporation spatially overlapping with damaged sites, which was significantly higher than signal identified in non-damaged sites in untreated samples (supplementary figure 6B). DM1 induced an increase in the number of damaged sites per microtubule (figure 6J) and abolished repair of damaged sites (figure 6K). This was rescued by the addition of NF1, which led to a strong increase in the fraction of repaired microtubules, with or without DM1 (figure 6J-K, supplementary figure 6A).

We conclude that NF1 enhances microtubule dynamics and enhances the repair of microtubule lattice damage. We also partially revise the mechanism of action of maytansinoids, which not only induce microtubule depolymerization from the extremities but can also cause microtubule lattice damage.

## Discussion

In this work we provide evidence for a RAS-independent role for NF1 in regulating microtubular dynamics and integrity. Its loss impacts chromosomal stability but may be exploited as synthetic lethality with maytansinoid-based ADCs like T-DM1.

An interaction between microtubules and NF1 was first observed in the 90s ^18,34,35^, but its physiological significance has remained poorly characterized to date. We demonstrate that NF1 has all defining features of a *bona fide* MAP, as it co-sediments with microtubules in vitro, induces microtubule bundling, decreases the critical concentration for tubulin polymerization and enhances microtubule dynamic instability. NF1 is crucial for normal mitosis, since its loss leads to increased mitotic plate size, delayed mitosis exit, accumulation of supranumerary centrosomes and chromosome missegregation. These mitotic defects collectively favor chromosomal instability (CIN) that leads to low-grade aneuploidy, identifiable both in cell lines and in patients. NF1-induced CIN may contribute to both decreased sensitivity to antiblastic agents and the metastatic phenotype ^36,37^, explaining the frequent occurrence of NF1 as a sub-clonal, late-appearing genetic alteration during tumor evolution ^38^.

Importantly, NF1 effect on microtubule dynamics and mitosis appears independent from its activity on RAS. The expression of a constitutively active, oncogenic RAS mutant does not reproduce the sensitization to DM1 induced by NF1 ablation; furthermore, NF1 enhancement of microtubule dynamics occurs in vitro in the absence of any additional protein, and can therefore be defined an intrinsic property of NF1.

Two regions residing in distant areas of NF1 (a C-terminal domain and a tubulin-binding domain next to the RAS-interacting domain) have been proposed to mediate interaction with tubulin ^18,19^ but structural data are currently missing. A testable and fascinating hypothesis is that MT dynamics may be enhanced by NF1 by promoting the intrinsic GTP-hydrolizing activity of beta-tubulin, which favors the stabilization of the tubulin newly incorporated in the lattice. This model would recall the established GAP function of NF1 on RAS. Early experiments did not identify a GAP activity using tubulin as a substrate; tubulin was, however, able to decrease NF1 GAP activity on RAS ^17^, suggesting that the two GTPAses may be competing for the same active site on NF1. The structural and functional domains responsible for the MT-NF1 interaction remain to be characterized and are the subject of ongoing research.

We show that DM1 treatment induces intra-lattice discontinuities, which are repaired more efficiently in the presence of NF1. The concept that MT lattice bears physiological discontinuities that are constantly repaired in vivo is relatively new ^21^, and may inform a better understanding of the mechanism of action of maytansinoids. The maytansin binding site, uniquely among MT-targeting agents, resides at the interface between two tubulin dimers ^6^. In the current paradigm, maytansin is thought to be able to bind only free dimeric tubulin and microtubule ends, leading to microtubule depolymerization from the extremities. This model is supported by experiments specifically assessing either de novo tubulin polymerization or microtubule instability ^7,8^. Our TIRF experiments using stabilized microtubules allowed to show that DM1 treatment leads not only to microtubule shortening but also to intra-lattice reduction in tubulin density, which in extreme cases lead to full microtubule breakage. This is likely allowed by the exposure of intra-lattice maytansin binding sites due to the lattice discontinuities that are now known to be generated in in vitro-stabilized microtubules ^39^. An important consequence of this model is that the number of available maytansin binding sites is not only determined by the number of microtubule ends but also by the number of damage sites; therefore, cell sensitivity to maytansin is determined by the efficiency of damage rapair. We speculate that other pathophysiological manifestations of NF1 loss, such as neurodevelopmental defects observed in neurofibromatosis, may be revisited as due to microtubule dysfunction and not, as currently proposed, to altered RAS signaling.

NF1 status is associated with differential sensitivity to endocrine therapy ^11,40^, targeted therapy (ref ^12^ and this study) and ADCs (this study). Mutations that confer resistance to payload in a minority of patients have been identified for T-Dxd^41^, the other major ADC approved in BC, but being acquired de novo they are of limited clinical utility as they cannot be assessed to identify nonresponders a priori. NF1 deficiency represents, to our knowledge, the first ADC payload-associated predictive biomarker identified to date. If these results are confirmed in independent studies, NF1 status may prove particularly useful in situations in which the identification of strong responders to T-DM1 may lead to its prioritization over T-Dxd, a more effective but also more toxic drug, like in the adjuvant setting, explored by the ongoing Destiny Breast 05 trial (NCT04622319), in which toxicity considerations play a stronger role. NF1 is covered by most routine NGS diagnostic panels (although with some limitations^10^) and can be implemented in routine diagnostics to drive treatment choice for breast and other cancers under the framework of precision medicine.

## Acknowledgments

BAD, EM, AC, EB, NR, AP, MRI, are PhD students at the European School of Molecular Medicine (SEMM), Milan, Italy.

We would like to thank the Genomic, Cell Culture, Flow Cytometry, and Molecular Pathology facilities at IEO for their support, dr Marina Mapelli at IEO for constructive discussion and the gift of recombinant NuMa, and dr Fabrice Andre at GR for constructive discussion

## Funding

This project was funded by the following grants:

For LM:

-My First AIRC grant n 25791.

-Italian Ministry of Health, Ricerca Corrente 2020-2022 and Ricerca Corrente di Rete (ACCORD) 2022-2024s.

-European Union – Next Generation EU – PNRR M6C2 – Investimento 2.1 Valorizzazione e potenziamento della ricerca biomedica del SSN – Project Code: PNRR-MAD-2022-12376934 - Capofila Dr Pier Giuseppe Pelicci.

For SS:

-Italian Association for Cancer Research grant AIRC-MFAG 2018 - ID. 21665

-Italian Association for Cancer Research grant AIRC Bridge Grant 2023 - ID. 29228

-Italian Ministry of Health, Ricerca Finalizzata grant GR-2018-12367077

-Fondazione Cariplo

-Rita-Levi Montalcini program from MIUR

-Italian Ministry of Health with Ricerca Corrente and 5 × 1000 funds For ZL:

-project National Institute for Neurological Research (Programme EXCELES, ID LX22NPO5107) - Funded by the European Union - Next Generation EU, institutional support from CAS (RVO: 86652036) and Imaging Methods Core Facility at BIOCEV supported by the MEYS CR [LM2023050, Czech-BioImaging].

For AD

-Breast Cancer Alliance

BAD is the recipient of a FIEO fellowship 2023

MRI is supported by an AIRC Fellowship (ID 26738-2021)

## Author contribution

Conceptualization: BAD, EM and LM. Methodology: BAD, EM, GT, EB, AC, GF, GC, SM, CS, DK, NR, AP, MRI, ED, AF, CJ, EGR, SR, DT, SS, MB ZL, LS, PGP, LM. Verification: EM, GT, EB, AC. Formal Analysis: BAD, EM, GT, EB. Investigation: BAD, EM, GT, EB, AC, GF, GC, SM, CS, DK, NR, AP, MRI, EGD, EM, MM, AF, AD, EN, CR, CJ, EGR, SR, DT, AM, AD, PDA, SS, MC, MB, ZL, LS, PGP. Writing: BAD, EM, GT, AC, EB, LM. Writing review: all authors

## Competing Interests

All authors declare no conflict of interest in relation to the submitted work.

Outside the submitted work, EG-R declared relevant relationships (advisory fees, honoraria, travel accommodation and expenses, grants and/or non-financial support) with AstraZeneca, Exact Sciences, GSK, Novartis, MSD, Roche, Sophia Genetics, Thermo Fisher Scientific.

AD declared relevant relationships as Scientific Advisory Board: Pfizer, Biotheranostics and Speaking honoraria: OncLive

## Lead contact

Further information and requests for resources and reagents should be directed to and will be fulfilled by the lead contact, Luca Mazzarella.

## Materials availability

All unique/stable reagents generated in this study are available from the lead contact with a completed Materials Transfer Agreement

## Data and code availability

Publicly available data:

GENIE cbioportal webserver version 13.0 (https://genie.cbioportal.org/) selecting breast cancer sample (15210). We selected samples that had been sequenced with the following panels that contain the NF1 gene: MSK-IMPACT341, MSK-IMPACT410, MSK-IMPACT468, GRCC-MOSC3, UCSF-NIMV4-TN, UCSF-NIMV4-TO Segmentation data was downloaded from cBioPortal using ASCETS (Arm-level Somatic Copy-number Events in Targeted Sequencing. ASCETS was run using UCSC cytoband coordinates (reference build hg19) provided in the ASCETS github repository (https://github.com/beroukhim-lab/ascets/blob/master/cytoband_coordinates_hg19.txt). The RNA-seq dataset (fastq files, raw and normalized counts, and DEGs) generated during this study are publicly available at GSE267855.

Personalized DNA sequencing data cannot be shared for privacy

The custom computer code generated for this project are publicly available through GitHub: https://github.com/mazzalab-ieo/tol_NF1_scripts.

## Materials & Methods

### Cell lines and cell culture

BT-474, SK-BR-3, HCC1954 and MDA-MB-231 cells were purchased from American Type Culture Collection (ATCC). The two former were cultured in Dulbecco’s modified Eagle’s medium (DMEM, 12800-017; Gibco, Grand Island, NY, USA) while the latter in RPMI-1640 (GE healthcare, Memphis, TN, USA). Both were supplemented with 10% fetal bovine serum (FBS, 10099-141; Life Technologies, Carlsbad, CA, USA), 100 U/mL penicillin and 100 mg/mL streptomycin and kept in a humidified 5% CO_2_ atmosphere at 37°C.

Cell lines are periodically tested for authentication using the ProMega geneprint 10 PCR-based kit (cat n B9510). Cells are never kept in culture for longer than a month or 10 passages, whichever is shortest.

Cell lines are periodically tested and for Mycoplasma contamination via the Uphof/Drexler protocol at every new restock, on average every 3 months.

### CRISPR-Cas9 generation of NF1KO clones

A guide oligonucleotide (sgN1, FWD: ACGGCCTGGACCCATTCCAC; REV: GTGGAATGGGTCCAGGCCGT) targeting exon 1 was selected based on best on-target/off-target scores provided by Benchling (https://benchling.com/), San Francisco, CA, USA. The guide was cloned into pSpCas9(BB) 2A-GFP (PX458) plasmid421, a gift from Feng Zhang (Addgene #48138). A DNA fragment containing 248nt of the NF1 coding region of interest was PCR-amplified from single-strand cDNA obtained from parental and clonal cells using QIAquick Gel Extraction Kit (QIAGEN). Incorporation was confirmed by Sanger sequencing the amplicon. All three cell lines were subsequently transfected with the plasmid containing the NF1 gRNA in parallel with empty vector controls using Lipofectamine 2000 according to manufacturer’s instructions. Seventy-two hours later, GFP-positive cells were single-cell sorted in 96-well plates. For the ribonucleoprotein (RNP)-mediated knockout of NF1 performed on BT-474 cells, Gene Knockout Kit v2 was ordered from Synthego (https://www.synthego.com/) and the Cas9 protein was purified in the institute, as a gift of L Rizzuti (Department of Experimental Oncology, European Institute of Oncology, Milan). RNP were assembled and lipofected into cells following the manufacturer’s instruction. After clonal expansion, knockout efficiency was analysed by western blot against neurofibromin and further characterised with NGS.

### Clinical study

Data were combined and shared under a data use agreement and approved by the institutional review boards (IRB) of the five sites (Washington University School of Medicine St. Louis, MO; IRB#202101147, Northwestern University, Chicago, IL; IRB#STU00214133, Massachusetts General Hospital, Boston, MA; IRB#2013P000848 and Weill Cornell Medicine/New York Presbyterian, New York, NY; IRB#1812019858, European Institute of Oncology, UID 4232). Data from Washington University, Northwestern University, Massachussets General Hospital and Weill Cornell were extracted from the combined database of the PMAC, precision medicine academic consortium.

The requirement for informed consent was waived by the IRB for this deidentified analysis. The study was performed in concordance with the Health Insurance Portability and Accountability Act and the Declaration of Helsinki.

Criteria for patient inclusion were:

-history of histologically confirmed, metastatic HER2+ BC

-treatment with T-DM1 in the 2^nd^ or 3^rd^ line in the metastatic setting. Therapy lines received in the neoadjuvant/adjuvant setting were not included in the computation

-no prior treatment with another ADC

-no concomitant treatment with another anticancer therapy, including endocrine therapy

-availability of an NGS assay with a panel covering NF1 performed on a relevant sample obtained after the diagnosis of metastatic stage. NGS assays included either FDA-approved liquid biopsy assays (Foundation 1 Liquid and Guardant360) or WES (described below). To preserve privacy, only data relative to NF1 were shared among collaborators.

The patients were independently extracted from each database, as follows:

-IEO: 111 patients treated with T-DM1 --> 79 patients excluded for lacking metastatic tissue sample or low amount/quality DNA after extraction è 32 patients left

-GR: 78 patients with mBC and available NGS --> 67 patients excluded for not having been treated with T-DM1 è 11 patients left

-PMAC: 1580 patients with mBC and available NGS --> 1537 excluded for not having been treated with T-DM1 è 43 patients left

PMAC, GR and IEO dataset were pooled in a single database (n=86) for the further check of homogeneity of inclusion criteria. 52 patients were discarded because of one of i) T-DM1 not administered in 2^nd^/3^rd^ line, or

ii) prior administration of another ADC, or iii) concomitant treatment with another anticancer therapy, including endocrine therapy

Mutations were analysed by alpha missense^42^ and RENOVO^43^ and considered to be pathogenic on the basis of at least one of the two algorithms.

Basic demographic characteristics are shown in supplementary table 1, NF1 mutations in supplementary table 2.

The efficacy endpoint was Progression-Free-Survival (PFS), defined as the time between the date of T-DM1 start and the date of the first clinically confirmed progression or death. Sample size was not prespecified given the exploratory and real-world nature of the study and the lack of preliminary data for differential response. PFS was modelled using the Kaplan-Meier method and log-rank test. No adjustment for covariates (which were only available for some patients) was performed.

### High-throughput drug screen

BT-474 WT and BT-474 KO cells were seeded (1 x 10^4^) in 96-well, white-bottom plates (Corning), in triplicates, and let grown overnight. Then, specified compounds were added in 9 dilutions plus their respective vehicle. After 96h, BrdU reagent was incorporated for additional 24h followed by fixation and detection according to manufacturer’s instructions (Cell Signalling Technology, #5492). Luminescence was read using a PHERAstar FSX Microplate Reader under a 425nm wavelength. The relative response was corrected compound-wise to the average vehicle response for each replicate. Data were analyzed in GraphPad Prism 9, and IC50 values were generated from best-fit curves. Data represent the mean ± SD.

Drugs were obtained from the following vendors: Doxorubucin (#44583), Paclitaxel (#T7402), Vinblastine (#V1377) from Merck, Carboplatin (#S1215), Pironetin (#CAY-28853) from Cayman Chemical, Tucatinib (#S8362), Lapatinib (#S1028), MMAE (#S7721) from Selleck. Pertuzumab, Trastuzumab, T-DM1 and T-Dxd were obtained from the IEO pharmacy.

### Growth curves

Cells were plated in triplicate at a density of 2×10^5^ cells/well for BT-474 and 1 × 10^5^ cells/well for HCC1954 in 12-well plates. After 24h, cells were treated with control (rituximab 10 μg/mL or DMSO), trastuzumab (10 μg/mL), T-DM1 (0.1 μg/mL), DM1 (5 nM) or T-DXd (0.1 μg/mL). For time point 0 (24h after seeding), and subsequently for all other time points (day 2, day 4, day 6, and occasionally day 8), cells were washed twice with PBS, trypsinized, counted using a hemocytometer and then treated again with fresh media containing the respective drugs.

### Colony formation assay

Duplicates of 6 x 10^2^ HCC1954 WT or HCC1954 WT and 1,2 x 10^3^ BT-474^WT^ or BT-474^KO^ cells were plated in 12-well-plates and left undisturbed at 37°, 5%CO2 for 4 days. Then, cells were treated with vehicle (rituximab 10 μg/mL or DMSO), Trastuzumab, T-DM1, DM1 or T-Dxd at the indicated concentrations for 21 days, with weekly change of spent media. At the end of treatment, cells were washed twice with ice-cold PBS and fixed with ice-cold methanol for 15 min. Afterwards, 0.5% crystal violet solution (Sigma V5265) was added to the plates and incubated at room temperature for 15 min. Plates were then washed with ddH2O, until the unbound crystal violet was removed and plates were dried at room temperature. Images were processed using the ColonyArea ImageJ plugin250.

### RasG12V overexpression and active Ras pull-down

KRAS^G12V^ from pDONR223_KRAS_p.G12V (Addgene #81665) was cloned on the lentiviral backbone pLenti CMV hygro dest (Addgene #17454). HEK293T cell transfection and infection of target cells was performed by Calcium Chloride followed by hygromycin B (InvivoGen) 0.5 mg/mL selection for 8 days.

Levels of active Ras were determined using the Active Ras Detection Kit (#8821, Cell Signalling Technology). Briefly, cell lysates (500 µL at 1 mg/mL) were treated in vitro with GTPγS or GDP to activate or inactivate Ras, respectively. The lysates were then incubated with glutathione resin and GST-Raf1-RBD for 1h at 4°C, washed and the resulting RAS-RAF1 complexes eluted from the resin by boiling in 2X SDS sample buffer. Proteins were resolved by SDS-PAGE, transferred to nitrocellulose and western blot analysis (20 µL of the eluted samples) was performed using the anti-RAS antibody supplied by the manufacturer. An anti-mouse IgG, HRP-linked Antibody (#7076) was used as the secondary antibody.

### Competitive co-culture

#### Vectors

The Tet-Off-H2B-GFP lentiviral vector422 and the H2B-mCherry retroviral vector423 were kindly provided by N Roda (Department of Experimental Oncology, European Institute of Oncology, Milan) and used in the following co-culture experiments. Tet-Off-H2B-GFP lentiviral vector was transfected in HEK293T cells together with 3rd generation packaging plasmids pMD2.G, pRSV-REV, and pMDLg/pRRE; H2B-mCherry retroviral vector was transfected in Phoenix-AMPHO cells instead. Viral supernatants were collected 2-4 days post-transfection and filtered through 0.45 mm filters. Filtered supernatants were concentrated by ultracentrifugation at 22.000 rpm for 2h at 4°C with OptimaTM L-90K ultracentrifuge (Beckman Coulter) and stored at −80°C (never refrozen).

#### Competitive co-culture

HCC1954^WT^ cells were infected with Tet-Off-H2B-GFP lentiviral vector while HCC1954^KO^ cells were infected with H2B-mCherry retroviral vector. Multiplicity of infection (MOI) = 2 and 8 μg/mL of Polybrene (Sigma-Aldrich) were used in both cases. Infected cells were selected via fluorescence activated cell sorting (FACS) using a BD FACSAria II. Sorted cells were subsequently mixed 1:1 and allowed to grow for up to 5 days in the presence or absence of 0.1 μg/mL of T-DM1. At day 0 (plating), 3, and 5, co-cultures were investigated by FACS to determine the fraction of wt and NF1 KO cells. Doublets of cells and cells with abnormal morphology were excluded for the analysis.

### Mouse xenograft models

#### Engineering of Luc-tagged HCC1954 cells (HCC1954-Luc)

The pLenti CMV Puro LUC (w168-1) was purchased from Addgene (#17477) and used in all in vivo experiments. This 3rd generation lentiviral vector allows for the constitutive expression of firefly luciferase under the CMV promoter. The vector also contains genes that encode for puromycin and ampicillin resistance. Both HCC1954^WT^ and HCC1954^KO^ cells were transduced with the pLenti CMV Puro LUC reporter vector at MOI = 5 in the presence of 8 μg/mL of Polybrene (Sigma-Aldrich). Infected cells were selected by puromycin (2.5 μg/mL for 72h; Vinci-Biochem) and allowed to expand for additional 8 days post selection. *Tumor implantation*

Tumors were implanted in 9-week-old Non-Obese-Diabetic/Severe Combined Immunodeficiency(SCID)/ Interleukin 2 receptor γ (NSG) mice, acquired from Charles River Laboratories, by intra-fat pad orthotopic injection of 10^6^ HCC1954^WT-Luc^ on the 9th mammary gland – left; and 10^6^ HCC1954^KO-Luc^ on the 5th - right) resuspended in sterile PBS (Thermo Fisher Scientific) and pre-mixed at a 1:1 dilution with growth-factor-reduced Matrigel (Corning) in a total volume of 30 μL. Tumor volume was weekly assessed via bioluminescence imaging (BLI) (IVIS-Lumina, Perkin Elmer). On day 14, once tumors were palpable (≥4 mm3), mice received a single intravenous injection of 4 mg/kg T-DM1 or NaCl 0.9%. Mice were weighed when cells were implanted (Day 0) and then twice weekly during the study. Experiment was terminated after 35 days. Animal experiments were authorized by the Italian Ministry of Health (n° 679/2020-PR)

#### Bioluminescence imaging (BLI) acquisition and analysis

The images were acquired using PerkinElmer’s IVIS Lumina Series III instrument wavelengths (600-800 nm). Scans were taken with an integrated CCD camera (Andor, Belfast, UK) supercooled down to −80 °C, with a 25 mm focal length lens (Navitar, Rochester, NY). The camera pointed straight down and was focused 10 mm above the imaging membrane. The F-number was kept at f/0.95 throughout the study. Prior to imaging, 200 µL of D-Luciferin (10 mg/mL, XenoLight, Perkin Elmer) were injected intraperitoneally and mice were anesthetized in induction chambers with 1-4% isoflurane. With animals in the supine position, BLI images were acquired after 10 min after luciferin injection. An exposure time of 2s and binning of 4 was used at the beginning of the study, and imaging parameters were updated as the tumors became brighter throughout the study, in order to maximize sensitivity of bioluminescence and avoid pixel saturation. Tumor burden was represented as total flux (photons/s), which is average radiance (flux per unit area and unit solid angle) integrated over the region of interest, using the Living Image v4.7.3 in vivo software package (Perkin Elmer Inc).

#### Immunohistochemistry

Formalin fixed and paraffine embedded (FFPE) tumour sections of 3µm thickness were cut and left for an overnight incubation at 37°C before staining. All immunohistochemistry (IHC) were performed using Bond III IHC autostainer (Leica Biosystems). For a full automated IHC staining, the antibodies rabbit monoclonal anti-Caspase 3 Activated (#9661 - Cell Signaling, final dilution of 1:250), rabbit anti-Cleaved Caspase 7 (#9491 - Cell Signaling, final dilution 1:100), rabbit monoclonal anti-cleaved Parp (#5625 - Cell Signaling, final dilution 1:50), and mouse monoclonal antibody anti-BCL-2 (#ab692 - Abcam, final diluition 1:100) were unmasked with EDTA Ph9 (Bond Epitope Retrieval Sol2 Leica AR9640). All the antibodies were diluted with Bond Primary Antibody Diluent (AR9352 - Leica Biosystems), and BOND IHC Polymer Detection Kit (DS9800 - Leica Biosystems) was used to detect antibodies with DAB chromogen and counterstain tissues with haematoxylin. H&E staining was performed with automated Leica Biosystems ST 5020 instrument, using Kit Infinity 2000 Test (3801698-Leica Biosystems) and pictures of stained sections were acquired with the Aperio ScanScope XT instrument.

### Immunoblot analyses

Cells were lysed in RIPA buffer (150 mM NaCl; 1% NP-40; 0,5% Sodium deoxycholate, 0,1% SDS, 25mM Tris, pH 7.4 in ddH2O) supplemented with PhosSTOP (Roche) and protease inhibitors (Roche). A protein fraction was obtained by centrifuging at 10.000 × g for 20 min at 4 °C. Supernatant was collected and protein concentration was determined using the Pierce™ BCA protein assay kit following manufacturer’s instructions (Thermo Fisher Scientific, Massachusetts, USA). Each sample was separated by non-reducing SDS-PAGE with Precision Plus Protein Dual Color Standards (Bio-Rad) and transferred to nitrocellulose membrane overnight at 4 °C. Where applicable, western blots were cut horizontally to allow the detection of different proteins within a single experiment. Blots presented in figure panels were derived from the same experiment and processed in parallel. The rest of the antibodies utilized here are: rabbit anti-cleaved PARP (Asp214, D64E10, XP® 5625); mouse anti-Bcl-2 (100/D5); mouse anti-GAPDH (6C5); rabbit anti-β-Tubulin (9F3); mouse anti-vinculin (A250291). Donkey anti-mouse IgG or goat anti-rabbit IgG secondary antibody conjugated to horse-radish peroxidase (Jackson ImmunoResearch Laboratories, Inc., Pennsylvania, USA) were probed against primaries 45 minutes at room temperature and detected bands were visualized using ECL reagent (Novex™ ECL Chemiluminescent Substrate Reagent Kit, Thermo Fisher Scientific, Massachusetts, USA) and processed using Image LabTM software 6.0.1 (Bio-Rad).

### Flow cytometry

BrdU was pulsed for 1 h on BT-474 and SK-BR-3 cells (WT and NF1 KO) treated with T-DM1 or vehicle for 36 h. Cells were fixed and stained according to manufacturer’s instructions (FITC BrdU Flow Kit, BD Biosciences). Samples were manually loaded, with acquisition criteria of 10.000 events or 3 minutes for each tube. During acquisition preview, gates for cells were adjusted in the FSC-A *vs* SSC-A plot, and the DNA 7-AAD-A voltage was adjusted to place the mean of the singlet peak (G0/G1) at 50.000 in the histogram. Cell cycle gates were adjusted as needed to encompass the G0/G1, S, and G2/M populations. Data was analyzed using FlowJo x.10.0.7r2 software (Tree Star).

To assess the effect of T-DM1 treatment on cell cycle distribution, harvested cells were fixed with 70% cold ethanol (Panreac Applichem) in PBS (Thermo Fisher Scientific), and stored at 4°C. After 8h, cells were centrifuged and resuspended in PBS (Thermo Fisher Scientific). DAPI (Sigma-Aldrich) was then added at the final concentration of 50 µM overnight at 4°C. Cell cycle distribution was analysed by FACS and differences in cell cycle distribution were evaluated through the χ2-test.

For HER2 expression analysis, 2 x 105 MDA-MB-231, BT-474 WT and KO cells were pelleted at 3000 rpm for 5 minutes. Cells were blocked with 5% BSA in PBS for 30 minutes and then incubated with primary antibody anti-HER2 Alexa Fluor-488 conjugate (#FAB1129G-025, Biotechne) diluted in 1% BSA in PBS for 1 hour at 4° C in agitation. Finally, cells were washed with 1% BSA in PBS and resuspended in 2% BSA in PBS). Flow cytometry was performed on BD FACS Celesta and all data were analysed using FlowJo 10.

### Immunofluorescence and confocal microscopy

#### Internalisation assay

Three batches of 7.5 x 10^5^ BT-474 cells were seeded on 0.5% gelatin-coated coverslips and allowed to attach overnight. After a 15 min T-DM1 or T-DXd pulse (1.5 μg/mL), drugs were washed off and one batch of cells was immediately fixed with PFA 4% for 15 min RT and analysed for baseline membrane impregnation. The remaining batches received fresh warm media and were incubated at 37°C for 24 h, with or without chloroquine (CQ, 5 μM). Cells were then fixed as previously described and all three batches were permeabilised with Triton X-100 0.5% in PBS and a sheep polyclonal anti-Human IgG-Cy5 (AC112S, Sigma) was used for the detection of T-DM1 and T-DXd; a FITC-phalloidin (P5282, Sigma) was used for actin and nuclear counterstaining was done with DAPI. Images were acquired with an SP8 confocal microscope (Leica Microsystems GmbH, Wetzlar, Germany) and a 63x/1.4NA oil immersion objective lens. Multichannel, Z-stack images were acquired with a voxel size of 72×72×200 nm3 by 3 different PMT detectors. To determine internalisation, single cells were segmented and the mean cytosol fluorescence signal of each single cell was evaluated (n = 28).

#### Chromosome misalignment analysis and centriolar quantification

BT-474 WT and BT-474 KO cells were seeded as per internalization assay. After 24h, cells were synchronized in late G2 with RO3306 (Sigma-Aldrich) at 9 µM for 39 or 22.5h, respectively (half their doubling time) at 37°C. Cells were then washed three times with warm medium to release from G2 arrest and put back at 37°C. After 50 min, cells were treated with MG-132 (Tocris) 10 µM for 90 min at 37°C. Finally, cells were fixed either by immersion in cold methanol at −20°C for 4 min followed by rehydration in PBS for 10 min or with 4% paraformaldehyde (PFA) for 15 min followed by permeabilization with 0.5% Triton X-100 for 20 min. For double thymidine block (DTB), SK-BR-3 cells were grown to a 40% confluence instead and 2 mM of thymidine was added. After incubation at 37°C for 18h, cells were washed with PBS and put back in culture with fresh media for 9h. After that, thymidine was added again for another 15h. Cells were then washed three times with warm media for release and fixed after 8h with 4% paraformaldehyde. Fixed cells were all blocked with 3% BSA for 1h and incubated in primary antibodies diluted in PBS-Tween20 0.1%, 3% BSA and 0.02% NaN₃. Primary antibodies used for immunofluorescence analysis included mouse anti-a-tubulin (T5168), human anti-centromere protein (15-234), rabbit anti-phospho-histone H3(Ser10) (06-570), and rabbit anti-CEP63 (06-1292). Primary antibodies were detected using Alexa Fluor-labelled secondary antibodies (Life Technologies) while DAPI (D9542) was used for nucleic staining. Coverslips were mounted using Mowiol and images were acquired as previously reported. Cell phenotypes were scored visually by counting non-overlapping FoVs in a raster scan pattern across the coverslip. Eight-bit images were exported and figures were prepared using Affinity Designer (Serif, West Bridgford,UK). Image data were quantified using ImageJ, and graphical representations and statistical analyses were performed using Graphpad prism 9.1.1.

#### Chromosomal and microtubular dynamics during mitosis

BT-474^WT-H2B-GFP^ and BT-474^KO-H2B-mCherry^ cells were seeded (40 x 10^3^ cells) in µ-Slide 8-well glass bottom chambers (80827, Ibidi). Twenty-four hours later, growing cells were stained for 1.5h before acquisition with the SiR-tubulin probe (#SC002, Spirochrome; λabs 652 nm/λfl 674 nm) at 1 μM. Time-lapse microscopy was performed using an inverted microscope (Nikon Eclipse Ti) with a 20X/0.75NA objective. The microscope was equipped with an incubation chamber maintained at 37°C in an atmosphere of 5% CO2 and acquisitions were made every 4 min. A total of 28 cells were analysed for BT-474^WT-H2B-GFP^ and 38 for BT-474^KO-H2B-mCherry^.

#### Microtubule repair

HCC1954^WT^, HCC1954^KO^ and NF1 WT re-expressing cells were seeded (1,5 x 10^5^ in 12-well plates) on coverslips coated with 0,5% gelatin and allowed to attach overnight. One batch of cells, used as control, was fixed in methanol at −20° C for 3 minutes, followed by rehydration in PBS-Tween0,1%. Another batch of cells was treated with DM1 at 1uM for 45 minutes at 37°C and then fixed as indicated above. Fixed cells were then blocked with 5% Normal Donkey Serum (Jackson ImmunoResearch #017-000-121) + 2% BSA + 0,3% Triton X-100 + 0,1% Sodium Azide in PBS for 1 hour and subsequently incubated with primary antibody mouse anti-alpha Tubulin (SigmaAldrich, clone DM1A, #T9026) diluted 1:500 in IF buffer (1x TBS + 0,1% Triton X-100 + 2% BSA + 0,1% Sodium Azide in PBS) for 2 hours. Primary antibody was detected using Alexa Fluor-488 secondary antibody (Life Technologies) diluted 1:400 and incubated for 45 minutes at RT. DAPI (Sigma, D9542; 1:1000) was used for nucleic staining. Images were acquired with using a CrestOptics confocal spinning disk X-light V3 with a 100x/1.49 NA objective lens (121 FOV for each condition).

### Live cell imaging of FUCCI(Ca)-expressing cells

#### Model generation and data acquisition

Stable FUCCI-expressing model was generated by co-transfecting the pCSII-EF-MCS vector encoding the FUCCI(Ca) ^27^ probe with the packaging plasmid (pCAG-HIVgp) and the VSV-G-/Rev-expressing plasmid (pCMV-VSV-G-RSVRev) into 293T cells. High-titer viral solutions were prepared and used for transduction into BT-474^WT^ and BT-474^KO^ cells by two rounds of infection of 4 viral particles per cell (MOI 1) followed by flow cytometry-based sorting (FACSAria cell sorter, BD). For live cell imaging, 0.5 x 10^5^ cells were seeded in µ-Slide 8-well glass bottom chambers (80827, Ibidi). Twenty-four hours later, T-DM1 or control (rituximab) were added at the indicated concentrations 30 minutes before starting acquisition. Images were acquired with a Yokogawa Spinning Disk Field Scanning Confocal System equipped with 20X/0.5 dry objective for 90h.

#### Automated cell cycle/cell fate analyses

For each experimental condition, we tracked cells in the mCherry (Red,R) and mVenus (Green,G) channels using TrackMate 7.2.0 (Ershov et al., 2021; Tinevez et al., 2017) on nine different fields of view (FoV), for a maximum of 530 frames, 10 min apart from each other, over 90h. Only cells that remained alive for at least 12h were followed by the tracker. For those, intensities on both channels were measured for each time frame starting from time 0 (T0).

#### Pre-processing

We first removed cells showing channel intensities lower than 200 grey levels for all the observed time lapse. Then, we applied a sliding average (window width = 5 points) to reduce noise and to smooth intensity signaling.

#### Algorithm to identify cell cycle phase

Using the sliding average values of both channel intensities for a cell of interest as input, we identified the cell cycle phase (G1, S, G2/M) and the “Lost by the tracker” condition (“Lost”) for each time point in the observed time lapse.

The first step is the identification of preliminary G1 and S frames, considering the channel intensities: if Red(*x_j_*)>Green(*x_j_*), the frame *x_j_* at time point *j* is classified as G1, otherwise as S.

Integrating the behaviour of channel intensities with tracker movies, we observed that the G2/M phase corresponds to a sharp increase in the mCherry (R) channel intensity when the cell is still in the S phase, matching to a decrease in the mVenus (G) channel intensity. The G2/M phase stops with a sharp decrease of the G channel intensity coupled with an increase in the R channel intensity, which determines the G1 entry. We thus set several rules to identify G2/M frames, studying the derivatives of red and green intensities for each frame x_i_ (d_red_(x_i_) and d_green_(x_i_)).

We classified as G2/M the frames x_j_ where:

- d_red_(x_i_) >1 and x_j_ was previously classified as S. To avoid the identification of small fluctuation as G2/M, we selected only groups of consecutive frames longer than 6 (equivalent to an hour) or, if none of those sequence is available, we selected only the longest sequence of consecutive frames;
- d_red_(x_i-1_)<1 and *j* is the last available time point;
- d_green_(x_i_) < (−5) and x_i_ was previously classified as G1;
- d_green_(x_i_) < (−1) with x_i_ previously classified as S and *j* > *j* where *j* = max*_k_ x_k_* such that *x_k_* is classified as G2/M
- both Red(*x_j_*) and green(*x_j_*) were higher than 200

After that, each frame is classified as G2/M, S or G1. We then checked frame by frame whether the order of phases in the cell cycle (G1 then S then G2/M) was respected. Residual misclassifications were further fixed with the following set of rules:

- in G2/M-S-S sequences, the first S is replaced by G2/M
- in G2/M-S-G2/M sequences, S is replaced by G2/M
- in G1-G2/M sequences, g2/m is replaced by G1
- in G1-S-G1sequences, S is replaced by G1

We then identified frames lost by the tracker, considering fluctuations in the ratio between the sliding averages of mCherry and mVenus channel intensities. This value should remain constant for dying cells: a coefficient of variation (CV) >0.1 on a sliding average (window width=7 points) was used to identify fluctuations.

We identified the last frame *x_m_* with CV>0.1: all its consecutive frames (*x_j_*, j>m) with a percentage of Red intensity 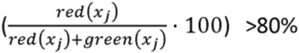 and dred(*x_j_*)<0.5 were identified as Lost, unless they had red(*x_j_*)≥200 or unless they were preceded by a S frame with increasing dred(*x_j_*).

We also set as Lost the last available frame if the previous one was defined as Lost.

Finally, to remove fluctuations, spurious “Lost” frames were reclassified as in the first part of the algorithm, while we forced spurious not “Lost” frames in assessed “Lost” sequences to the “Lost” class.

#### Analysis of identified cell cycle phases

we applied our algorithm to all the available FoV for each condition. Once we assigned a phase to each frame, we dissected the complete cell cycles identified. To plot the cell phases history we excluded cells whose first frame was classified as G2/M, in order to only include cells whose G2/M phase could be monitored from beginning to end. We compared the percentage of frames assigned to the G2/M phase in NF1 KO cells (vehicle, T-DM1 0.1µg/mL and T-DM1 1µg/mL) with the corresponding percentage in the WT population. We performed a Wilcoxon test for statistical analysis, with significance at 0.05.

We also considered the frequency of cell transitions between different phases and compared them across conditions with a Chi-squared test.

### Microtubule tip tracker EB3-GFP

HCC1954 cells were maintained in complete phenol red-free RPMI-1640 medium. Cells were subcultured onto µ-slide 8 well (ibidi Cat. No. 80826) and transfection was performed with LentiBrite™ EB3-GFP Lentiviral Biosensor (Sigma, Cat. No. 17-10208) according to the manufacturer’s protocol. The culture medium was replaced 24 h post-transfection. Fresh media containing mertansine 600pM or DMSO were added 48 h after transfection and 30 min before acquisition. Fluorescent microscopy was performed on a Nikon CSU-W1 Yokogawa Spinning Disk Field Scanning Confocal System equipped with a heating incubator chamber. Imaging was performed on cells with an appropriate level of transfection without overexpression patterns. Image sequences were acquired with 100x/1.45NA oil immersion objective and the image pixel size was set as 0.136 x 0.136 x 1 m^3^. Time-lapses were recorded with a frame step of 1 s for 60 s, with an exposure time of 20 ms. The number of individual cells for each experimental day was about 10. The EB3 fluorescence signal motion was analysed with the Fiji’s plugin TrackMate v. 7.12.1.

### CETSA melting curves

The cellular thermal shift assay was performed as previously described ^46^. Growing BT-474 cells were harvested, washed once with PBS, and diluted in Hanks’ Balanced Salt Solution (HBSS, Thermo Fisher Scientific) supplemented with PhosSTOP (Roche) and protease inhibitors (Roche) to 4 x 10^6^ cells/mL. The cell suspension was then mixed with DMSO or DM1 1 μM and kept on a wheel at 37°C for 2.5 h before aliquoting into PCR-tubes. Melting curves were generated by heating samples for 3 min in a Veriti Thermal cycler (Applied Biosystems) at the following stepwise temperature range: 57°C, 59°C, 61°C, 63°C, 65°C, 67°C. Cells were subsequently lysed with three rounds of freeze-thaw by alternating exposure of the samples to liquid nitrogen and 20°C in a PCR-machine. Samples were then resolved by SDS-PAGE gel, transferred to nitrocellulose and analyzed by immunoblotting against b-tubulin (9F3). For the quantitative analysis, we first normalized mean band intensities obtained for temperatures within our range on the mean band intensity obtained at the starting temperature (57°C), considering the treatment (DMSO or DM1). To understand which factors affected most the differences observed, we performed a two-way ANOVA separately for BT-474^WT^ and BT-474^KO^ normalized data. We considered as factors the treatment, the temperature and their combination.

### NF1 expression and *in vitro* purification

The NF1 isoform 2 recombinant protein was produced in insect cells starting from the plasmid R702-X38-635 encoding for His6-Hs.NF1opt (1-2839) (a gift from Dominic Esposito; http://n2t.net/addgene:159576; RRID:Addgene_159576). The vector was further modified by mutagenesis (GeneScript) by inserting a Prescission cleavage site between the His-tag and the NF1 coding sequence.

The protein was expressed by infection of High Five Cells (BTI-TN-5B1-4) (ThermoFischer Scientific) with the baculovirus obtained from the above-mentioned vector. Cell pellets were resuspended in lysis buffer (25 mM Hepes pH 7.5, 500 mM NaCl, 5% Glycerol, 1 mM DTT) supplemented with protease inhibitors cocktail Set III (Calbiochem) and DNAseI (PanReac AppliChem), lysed mechanically by dounce homogenizer and cleared by centrifugation at 70000 g for 45 min at 4°C. The supernatant was filtered through 0.22 µm filter (sartorius) and then loaded on a 5 ml HisTrap HP column (Cytiva) using an AKTA chromatography system. The column was washed with a buffer containing 38 mM imidazole and the protein was then eluted with a buffer containing 250 mM. The resulting sample was incubated overnight at 4°C with recombinant Prescission protease at a protein:protease ratio of 100:1 (w/w), to cut the His-tag. After cleavage, the protein was concentrated and loaded on a Superose 6 column (Cytiva) pre-equilibrated in SEC buffer (25 mM Hepes pH 7.5, 300 mM NaCl, 5% Glycerol, 1 mM DTT). Fractions containing monodisperse protein were concentrated in a 100 kDa cut-off Amicon ultra centrifugal filters (Millipore) and flash frozen in liquid nitrogen for further assays. Protein concentration was determined by measuring sample absorbance at 280nm using the extinction coefficient calculated from the amino acid sequence.

#### NF1 WT-rescue experiment under KO background

The pLVX-puro vector was kindly provided by A Polazzi (Department of Experimental Oncology, European Institute of Oncology, Milan) and was further modified by mutagenesis (GeneScript) by inserting the NF1 coding sequence. The pLVX-puro-NF1 lentiviral vector was transfected in HEK293T cells by Calcium Chloride. Viral supernatants were collected 2 days post-transfection and filtered through 0.45 mm filters. HCC1954 NF1 KO cells were infected in the presence of 8 μg/mL Polybrene (Sigma-Aldrich). Infected cells were then selected by Puromycin (4 μg/mL for 5 days; Vinci-Biochem) and allowed to expand for additional 5 days post selection. NF1 re-expression in HCC1954^KO^ cells was confirmed by immunoblot analyses.

### Mass photometry

NF1 molecular mass was measured at a final concentration of 10nM in 25mM Hepes pH7,5, 300mM NaCl, 5% Glycerol, 1mM DTT with the TwoMP instrument (Refeyn). The calibration curve was generated using the two peaks corresponding to 66 kDa and 132 kDa resulting from a BSA measurement.

### GAP-stimulated GTPase activity assay

We tested the functionality of our purified NF1 by evaluating its RAS-GAP activity through the GTPase-Glo™ assay (Promega), a bioluminescent assay that quantitates the amount of GTP remaining after a GTPase reaction. Reactions were assembled with 2μM Ras and 1μM NF1 in GTPase/GAP Buffer provided by the kit. Reactions were initiated by adding 5μl of 10μM GTP in GTPase/GAP Buffer containing 1mM DTT. The final reaction volume was 10μl. Reactions were incubated for 90 minutes at room temperature. To the completed GTPase reactions, 10μl of GTPase-GloTM Reagent was added and reactions were incubated for 30 minutes at room temperature. Twenty microliters of Detection Reagent were added, plates were incubated for 5–10 minutes at room temperature and luminescence was recorded using the GloMax®-Multi+ Detection System.

### Co-sedimentation assay

20µM Tubulin (Cytoskeleton Inc) was polymerized into stable Microtubules in General Tubulin Buffer (80 mM PIPES pH 6.8, 1 mM MgCl2, 1 mM EGTA), supplemented with 1 mM GTP and 50 µM Paclitaxel at 37°C for 20 minutes. For Microtubule-binding reactions, microtubules were diluted to a final concentration of 9µM in General Tubulin Buffer supplemented with 1 mM GTP, 50µM Paclitaxel and 60mM NaCl. 0,6 µM NF1 and 5 µM NuMA (a gift from Marina Mapelli, Department of Experimental Oncology, European Institute of Oncology, Milan) proteins were added to a final volume of 50µl. Reactions were incubated at room temperature for 15 minutes, transferred onto 100 µl of cushion buffer (80 mM PIPES pH 6.8, 1 mM MgCl2, 1 mM EGTA, 50 µM Paclitaxel, 50 % glycerol) and ultracentrifuged for 15 minutes at 400,000 g at 25 °C in a Beckman TLA100 rotor. The supernatant (50µl) was transferred from each centrifuge tube in SDS loading buffer. The cushion buffer was discarded very gently. The pellet was resuspended in SDS loading buffer 1x. Pellets and supernatants were analysed by SDS-PAGE.

### Turbidity assay

3 mg/mL Tubulin was mixed on ice in a 96 well plate with 0,6 µM NF1 or 10 µM Paclitaxel (positive control) or 10µM Nocodazole (negative control) in a final volume of 100μL in Tubulin Polymerization Buffer (80 mM PIPES pH 6.9, 2mM MgCl2, 0.5mM EGTA, 1mM GTP, 10% glycerol). The polymerization reaction was started by the increase in temperature from 4°C to 37°C upon transfer of the reaction to pre-warmed microtiter spectrophotometer. Tubulin assembly was monitored for one hour at 340 nm in kinetic mode of 61 cycles for each sample and the readings were recorded at an interval of 1 min (Infinite 200 PRO-Tecan plate reader).

### *In vitro* microtubule dynamics assays

Biotin-labeled tubulin as well as HiLyte647-labeled tubulin and TAMRA Rhodamine labeled tubulin were purchased from Cytoskeleton Inc. (T333P, TL670M and TL590M respectively).

GMPCPP-lattice microtubules (GMPCPP polymerized) were polymerized from 4 mg/ml HiLyte-647-labeled biotinylated pig-brain tubulin (12% biotin tubulin, 20% HyLite647-labeled tubulin, 68% unlabeled tubulin) for 1h at 37֯C in BRB80 supplemented with 2.7 mM GMPCPP (Jena Bioscience, NU-405). The polymerized microtubules were centrifuged for 30 min at 18000 x g in a Microfuge 18 Centrifuge (Beckman Coulter). After centrifugation the pellet was resuspended in BRB80.

### TIRF microscopy

Total internal reflection fluorescence (TIRF) microscopy experiments were performed on an inverted microscope (Nikon Eclipse Ti-E) equipped with 60x oil immersion objectives (Apo TIRF, Nikon) and Hamamatsu Orca Flash 4.0 CMOS camera. An additional 1.5x magnifying tube lens was used. Hilyte-647-labeled biotinylated microtubules and TAMRA Rhodamine labelled tubulin were visualized sequentially by switching between microscope filter cubes for Cy5 and mCherry channels. The imaging setup was controlled by NIS Elements software (Nikon). Flow chambers for TIRF imaging assays were prepared as described previously ^47^. Channels were treated with anti-biotin antibody solution (Sigma, B3640, 1 mg/ml in PBS) after 5 minutes, followed by one-hour incubation with 1% Pluronic F127 (Sigma, P2443).

#### Microtubule dynamics measurements

To determine how microtubules grow in the presence of NF1 or DM1 and DM1 with NF1 together we employed microtubule dynamic assay. We first mobilized microtubule Hilyte-647-labelled biotinylated microtubules and subsequently flushed in the assay buffer (BRB80, 1 mM GTP, 0.2% Tween20, 0.5 mg/ml Casein, 20 mM D-glucose, 0.22 mg/ml glucose oxidase and 20 ug/ml catalase) with free 20 μM HyLite-647 tubulin in the absence or presence of 100 nM NF1, 300 nM NF1 and/or 1 μM DM1. Time-lapse image sequences were recorded for 20 min at the rate of 1 frame per 10 s with an exposure time 100 ms. Data from three independent experiments was collected, each experiment was repeated at least on three days.

Microscopy data were analyzed using ImageJ (FIJI). Dynamic parameters of microtubules in the presence of NF1/ DM1/ NF1 with DM1. Dynamic instability parameters microtubules as polymerization rate, depolymerization rate, frequency of catastrophes and frequency of rescues was analysed using Kymographs, which were generated using KymographBuilder plug-in In FIJI by manually fitting lines along the microtubule tips.

#### Microtubule depolymerization assay

To determine if NF1 could rescue microtubule depolymerization we employed microtubule depolymerization assay in the presence of NF1 and/or DM1. First, we mobilized Taxol-stabilized TAMRA Rhodamine labeled biotinylated microtubules and subsequently washed them with wash buffer (BRB80, 0.2% Tween20, 0.5 mg/ml Casein, 20 mM D-glucose, 0.22 mg/ml glucose oxidase and 20 ug/ml catalase). Time-lapse image sequences were immediately started and the sample was imaged for 15 min at the rate of 1 frame per 5 s with an exposure time 100 ms. During the imaging, first microtubules were incubated in the washing buffer (without taxol) for 1.5 min. Subsequently microtubules were incubated in the assay buffer (BRB80, 0,1% methylcellulose, 1 mM GTP, 0.2% Tween20, 0.5 mg/ml Casein, 20 mM D-glucose, 0.22 mg/ml glucose oxidase and 20 ug/ml catalase) supplemented with 2 uM Hilyte-647-labeled tubulin (to slow down microtubules disassembly) in the absence or presence of 100 nM NF1/1 μM DM1/ 100 nM NF1 with 1 μM DM1 for 10 min. In the last step microtubules were washed with the wash buffer supplemented with 25% glycerol to prevent further lattice depolymerization.

### Microtubule damaged sites and intratubular tubulin repair assays

*In vitro* microtubule repair experiments were performed as described previously ^23^. Briefly, TAMRA Rhodamine labeled tubulin polymerized in the presence of 50 µM Taxol were first incubated without Taxol and tubulin for 1.5 min and subsequently with 5 μM HiLyte Fluor-488-labeled tubulin with or without 300 nM NF1 and/or 1 μM DM1 to promote repair. After 20 min, the residual free green tubulin was washed out with wash buffer (80 mM PIPES, 4 mM MgCl2, 1 mM EGTA, 0.5 mg/ml κ-casein and oxygen scavenger mix (50 mM glucose, 400 μg/ml glucose-oxidase, 200 μg/ml catalase, and 4 mM DTT)) supplemented with 25% glycerol to prevent microtubule depolymerization, in order to better visualize incorporation of green tubulin. Images were the acquired on an inverted TIRF microscope (Nikon Eclipse Ti-E) equipped with 100x oil immersion objectives (Apo TIRF, Nikon) and Hamamatsu Orca Flash 4.0 CMOS camera.

### Intensity quantifications along the lattice at damage site

#### Preprocessing

We obtained microtubule and tubulin signal intensities from two different experiments, for a total of 174 microtubules (86 and 88 respectively for experiment and 2). Microtubules were assigned to four conditions: control (22 for experiment 1, 27 for experiment 2); addition of DM1 (21 and 16); addition of NF1 (23 and 21); and addition of both NF1 and DM1 (20 and 24).

We first normalized each microtubule intensity profile I(x):

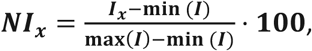

with respect to the maximum and minimum intensity along the whole profile (NI values between 0 and 100). We then computed the rolling mean of the NI (window=2), value used in the following analysis We used the rolling mean with a window of 2 also for tubulin intensity.

Our analysis focuses on microtubule shaft, not taking into account regions around microtubule tips. Those regions were identified by selecting the first and last peaks of tubulin intensity (R package pracma, https://cran.r-project.org/web/packages/pracma: selected patterns show at least 3 points with a decreasing tubulin intensity for the first peak and at least 3 points of increasing tubulin intensity for the last one). Microtubules with shaft shorter than 1.5 microns were removed from the analysis (Experiment 2: 3 for control and 1 for NF1+DM1 condition).

#### Damage sites identification

We first identified damage sites along the microtubules shaft, defined as microtubule regions long at least 0.5 micron and with an intensity lower than a certain threshold. The threshold is not uniquely defined along the microtubule, but defined point by point with an “adaptive” algorithm. First, the microtubule intensity profile is divided in ***m*** segments, according to ***m-1*** possible changes in the profile trend. Points where a change in trend occur are identified by the means of R package strucchange (https://cran.r-project.org/web/packages/strucchange). On each segment of the microtubule profile, we then fit a linear model, with the normalized intensity of the microtubule as dependent variable and the microns from the beginning of the segment as independent variable:

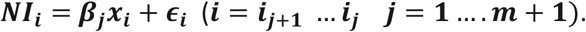

The threshold to identify damaged regions in each segment ***k*** is then defined as: **max(β*_k_***, ***mean***(*NI*)), where ***mean***(***NI***) is the average normalized intensity along the microtubule shaft. Regions below this threshold are considered damaged. To avoid considering sites with a too high intensity as damaged, we set the maximum of the threshold to 70.

#### Tubulin peaks identification

To identify regions with tubulin intensity peaks, we first set a threshold to remove background tubulin. Considering only microtubules in the control case we analyzed the maximum value reached by tubulin on the tips, which should describe, in terms of tubulin, the total intensity of the microtubule. Tubulin with intensity lower than 20% of the median of those values (220.69 for Experiment 1; 1000.29 for Experiment 2) was considered as background tubulin.

For each condition we then computed tubulin intensity quartiles along the body of the microtubules. To identify tubulin peak regions, we called peaks over two different runs. On each microtubule we searched for patterns with at least 4 increasing intensities before the possible peak and at least 4 decreasing intensities after it (R package pracma). In the first run we selected positions with the required and with a minimum height higher than the 3rd tubulin quartile in the specific condition; in the second set the 2nd quartile as minimum height. To consider only the significant contribution of the tubulin, “peak regions” were then identified as part of the peaks lying over the 2nd quartile in the first run and over the 1st quartile in the second run. The second run had an impact on the number of called peaks only when NF1 was added (NF1 and NF1+DM1 cases). To avoid background fluctuations, we removed peaks exceeding the background threshold defined above.

#### Damaged sites and intratubular repair analysis

We identified areas of intratubular repair by overlapping damaged sites with the regions of tubulin peaks. We thus studied the average level of tubulin in damaged vs non-damaged regions on the control case (Wilcoxon paired test), and applied permutation test to analyze differences in the number of damaged sites and of intratubular repairs across the four conditions.

### DNA sequencing

Genomic DNA extraction was performed using the DNeasy Blood & Tissue Kit (Qiagen), in accordance with the manufacturer’s instructions. DNA concentrations were carried out using Qubit dsDNA Broad Range quantification assay kit (Thermo Fisher Scientific).

For clone validation, libraries were generated using TruSight Rapid Capture kit in combination with the TruSight Cancer Sequencing Panel (Illumina) according to the manufacturer’s protocol. Briefly, 50 ng of gDNA was enzymatically fragmented and adaptor sequences were added to the ends. The tagmented DNA was amplified by PCR followed by purification. Target regions were captured with Cancer Sequencing Panel probes followed by PCR amplification and purification of the enriched library.

For WES, Target regions were captured with Twist Comprehensive Exome Panel probes followed by PCR amplification and purification of the enriched library.

Quantification of enriched libraries was performed with Qubit dsDNA High Sensitivity quantification assay kit (Thermo Fisher Scientific) and library size distribution was measured with Bioanalyzer 2100 and High Sensitivity DNA Kit (Agilent Technologies). Final DNA libraries sequencing was performed in Illumina NovaSeq 6000 platform using the NovaSeq 6000 S1 Reagent Kit 300 cycles (2 x 150 paired-end reads) (Illumina). Sequencing data were mapped against hg38 genome and analysed with Illumina DRAGEN Bio-IT Platform v4.0 using proprietary pipelines for variant calling. The resulting VCF with detected variants files were were annotated and classified with the GATK-Funcotator.

### RNA-seq

mRNA-seq libraries were prepared according to the TruSeq low sample protocol (Illumina, San Diego, CA, USA), starting with 1 µg of total RNA per sample. RNA-seq libraries were pair-end sequenced on an Illumina NovaSeq 6000 sequencing platform. RNA-seq data were mapped using the STAR aligner version 2.7.1a against the human genome (hg19). Read counts were calculated with htseq-counts with the standard UCSC reference GFF (https://hgdownload.soe.ucsc.edu/goldenPath/hg19/bigZips/genes/hg19.refGene.gtf.gz). Differential expression analysis was done with DESeq2 R package version 1.36.0. Genes of interest were selected using a false discovery rate (FDR) cut-off of 1×10^−4^

Functional enrichment analysis war performed using EnrichR R package version 3.1 on differentially expressed genes with log2foldchange < −1 in NF1 knock-out T-DM1 treated vs NF1 knock-out and log2foldchange > −1 and log2foldchange < 1 in wild-type T-DM1 treated vs wild-type. Queries were performed over four different molecular signatures: Gene Ontology Biological Process, Gene Ontology Molecular Function, Gene Ontology Cellular Component (https://www.gsea-msigdb.org/gsea/msigdb/download_file.jsp?filePath=/msigdb/release/2022.1.Hs/c5.go.v2022.1.Hs.symbols.gmt) and Hallmark (https://www.gsea-msigdb.org/gsea/msigdb/download_file.jsp?filePath=/msigdb/release/2022.1.Hs/h.all.v2022.1.Hs.symbols.gmt) (10.1073/pnas.0506580102, 10.1093/bioinformatics/btr260, 10.1016/j.cels.2015.12.004). Enriched terms were selected using a standard FDR cut-off of 1×10^−2^.

#### Arm-level Aneuploidy Assessment

GENIE data were retrieved from GENIE cbioportal webserver version 13.0 (https://genie.cbioportal.org/) selecting breast cancer sample (15210). We selected samples that had been sequenced with the following panels that contain the NF1 gene: MSK-IMPACT341, MSK-IMPACT410, MSK-IMPACT468, GRCC-MOSC3, UCSF-NIMV4-TN, UCSF-NIMV4-TO reducing the sample size to 6598 samples. Of this 58 are NF1 mutated. NF1 mutated samples were selected using the following criterium: samples with truncating mutations (frameshift insertion/deletion, non-sense mutations, canonical splice-site variants) or samples with missense mutations predicted “pathogenic” by the functional predictor AlphaMissense ^31^

Aneuploidy scores (AS) were quantified on segmentation data downloaded from cBioPortal using ASCETS (Arm-level Somatic Copy-number Events in Targeted Sequencing. ASCETS was run using UCSC cytoband coordinates (reference build hg19) provided in the ASCETS github repository (https://github.com/beroukhim-lab/ascets/blob/master/cytoband_coordinates_hg19.txt). AS was calculated with standard parameters with a minimum breadth of coverage of 0.5 and alteration threshold of 0.7.

### NF1 Mutated Patients Prevalence

To assess mutational prevalence of NF1 were selected two cohorts: i) GENIE (Genomics Evidence Neoplasia Information Exchange) cohort database (v13.0-public) and ii) TCGA PanCancer Genome Atlas (2018) Breast Cancer Cohort (10.1038/ng.2764).

Only patients with breast cancer were selected. Subsequently the cohorts were divided into two subcategories ERBB2 amplified and ERBB2 wilt-type.

TCGA PanCancer Genome Atlas Breast Cancer Cohort was used to select primary samples; GENIE cohort was used to selected metastatic samples.

Odds ratio were calculated as the ratio of gene frequencies in metastatic versus primary cohorts. Genes with less than 5% of mutated samples, considered poorly informative, were excluded to avoid false-negatives FDR overcorrection.

NF1 lollipop plots have been designed using ProteinPaint (https://pecan.stjude.cloud/proteinpaint).

### Statistical methods

Statistical differences on continuous dependent variables were based on two-tailed t-tests, after assessment of distribution normality via the Shapiro test. When at least one of the continuous variables was not normally distributed, significance was based on Wilcoxon test. Significance was considered at pvalue≤0.05.

In cases in which independent categorical variables had more than 2 values, we applied 1- or 2-way ANOVA, according to the number of independent categorical variables used and then applied Tukey’s HSD test to identify pairwise statistically significant differences. When time was one of the used variables, we instead performed a 3-way ANOVA with repeated measurements, followed by a post-hoc pairwise t-test. When the normality distribution hypothesis was not met, we applied Kruskal-Wallis with Dunn post-hoc test. In all those cases, given the small number of simultaneous hypotheses, a simple Bonferroni correction was performed (significance: adjusted pvalue<0.05).

Statistical differences in count distribution among categorical conditions were tested with Fisher’s exact test (variables with 2 factors), Chi-squared test in cases where the variables had more than 2 factors and Fisher-Pitman asymptotic permutation test when one of the categorical condition represented a numerical variable. Significance was considered at pvalue≤0.05.

Specific statistical tests for complex data are described in the relative method section.

---

**Supplementary Table 1.** Summary characteristics of patients analysed in figure 2 L-M

**Summary Table 2**: NF1 mutations considered for the retrospective clinical studied

**Supplementary figure 1.**
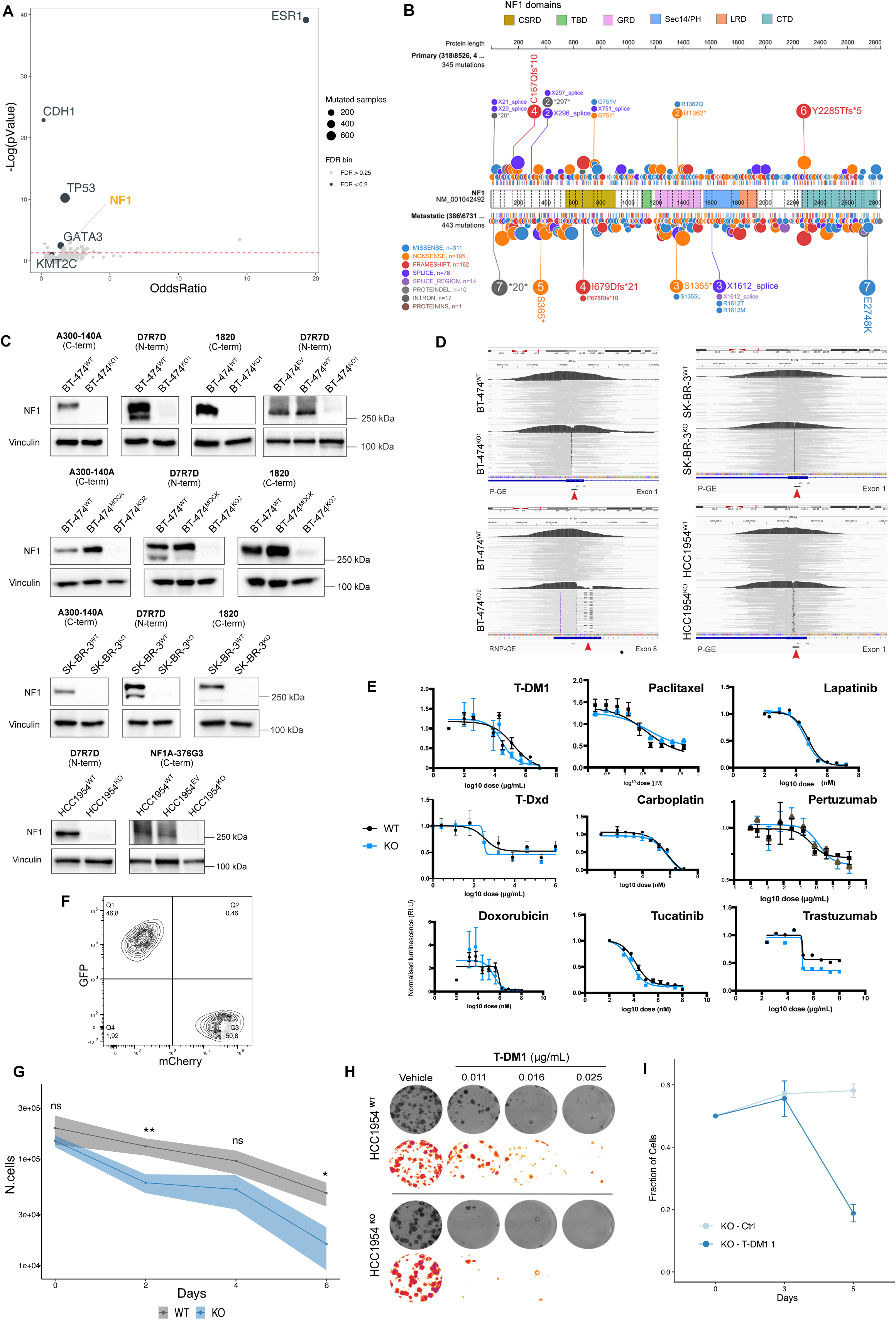
**A)** Comparison of mutation frequency in unmatched metastatic *vs* primary tumors in unsorted BC, from TCGA and AACR-GENIE. Plot depicts odds ratio (OR) *vs* -log p value of mutation frequency. **B)** Lollipop plot depicting frequency of point mutations in metastatic *vs* primary breast tumors in unsorted BC cohorts. **C)** Assessment of NF1 protein loss in multiple clones of CRISPR-Cas9-engineered BT-474, SK-BR3 and HCC1954 cells, using multiple NF1 antibodies. **D)** Representative IGV visualizations of the NF1 genomic region surrounding the guide RNA target site, in WT and Cas9-engineered clones. **E)** Dose-response curves derived from BrdU incorporation assays using nine therapeutic range concentrations for each compound among FDA/EMA approved agents used in the treatment of HER2+ BC. **F)** 8-days growth curves on HCC1954^WT^ and HCC1954^KO^ cells treated with T-DM1 0,1 μg/ml. **G)** Colony formation assay for HCC1954^WT^ and HCC1954^KO^ cell lines treated with T-DM1 at the indicated doses. **H)** Co-culture of HCC1954^WT-H2B-GFP^ or HCC1954^KO-H2B-mCherry^ cells; mean ± SD.

**Supplementary figure 2.**
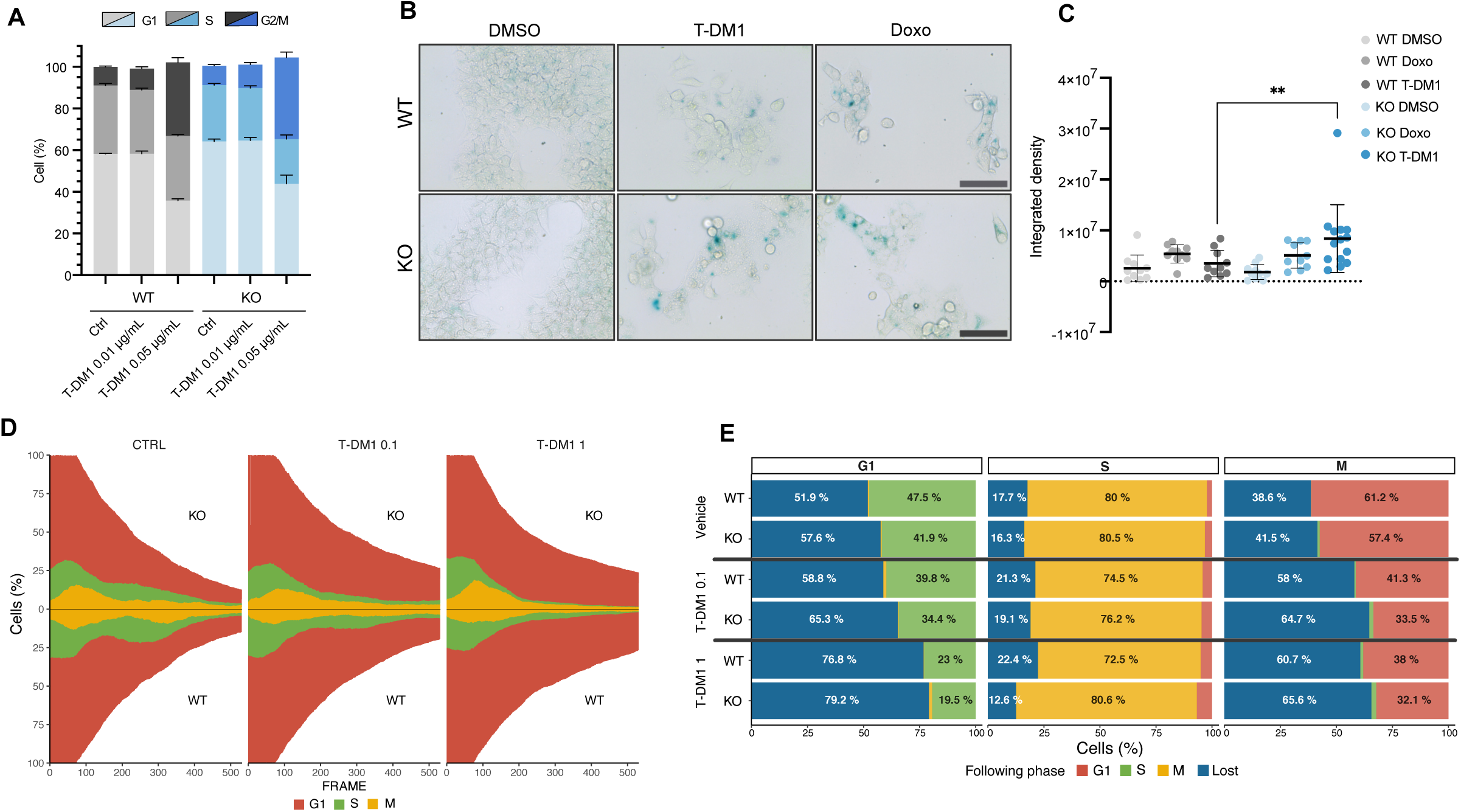
**A)** Propidium-iodide (PI) cell cycle analysis in SK-BR3^WT^ and SK-BR3^KO^ cells after 36h of T-DM1 at the indicated doses. Error bars are SD. **B)** Beta-galactosidase activity in BT-474^WT^ and BT-474^KO^ cells treated with vehicle (DMSO), T-DM1 (1 μg/ml) and Doxorubicin (200 nM) for 5 days (scale bar=1.5 mm). **C)** Quantification of panel A by integrated density (Mann Whitney U test, **p<0.01). **C)** Cell fate analysis of 200 cells/condition during 90h of T-DM1 exposure, showing a reduction in proliferating cells, longer G2/M arrest and higher G1 prevalence across time in BT-474^KO^ cells compared to BT-474^WT^. **E)** Phase transitions in FUCCI(Ca)-expressing cells displayed as percentages.

**Supplementary figure 3.**
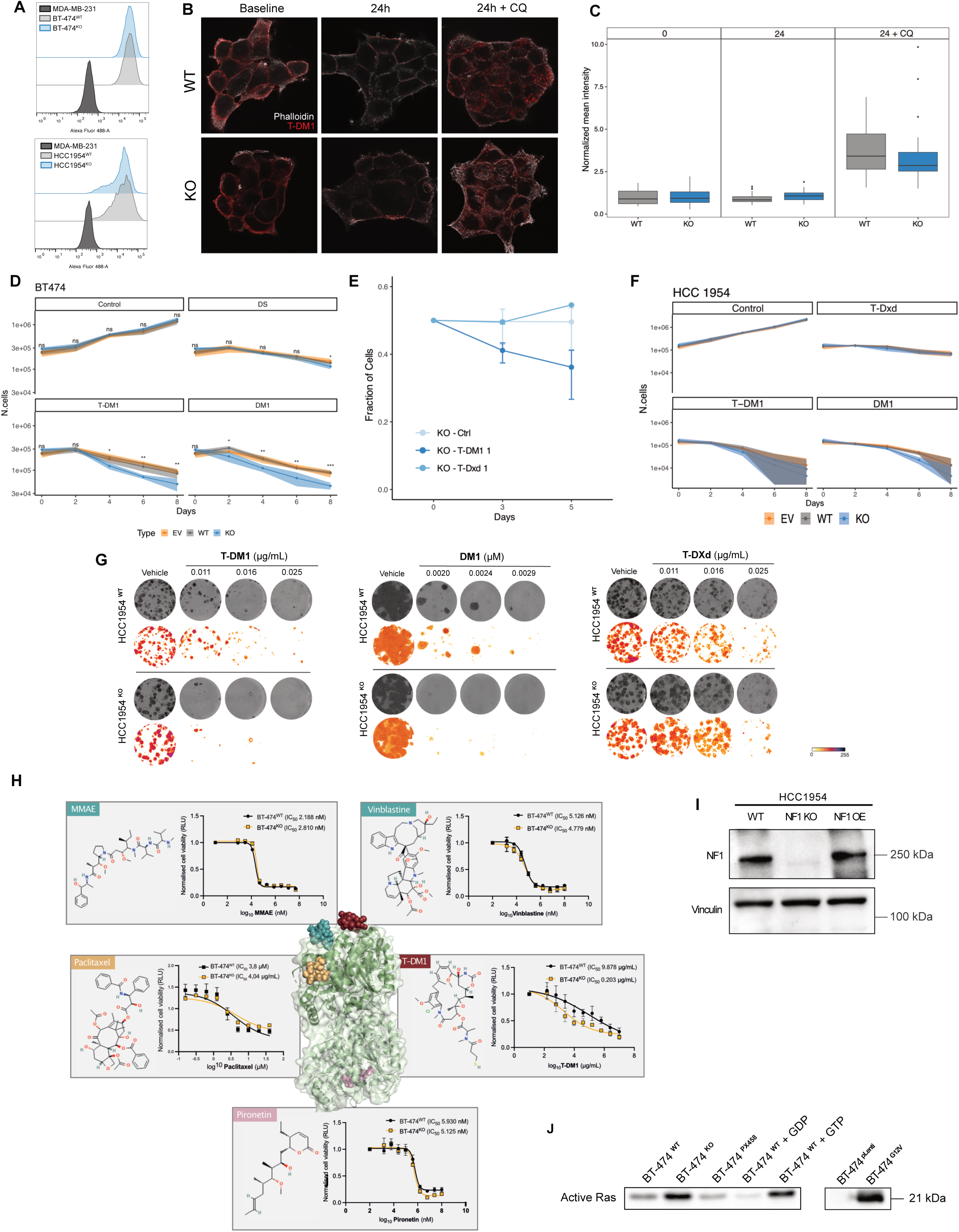
**A)** Assessment of HER2 expression in BT-474 and HCC1954 cells (WT and KO) by flow cytometry. MDA-MB-231 cells were used as negative control. **B)** T-DM1 internalization assay. BT-474^WT^ and BT-474^KO^ cells were incubated with vehicle or T-DM1 (1 μg/ml) for 24 hours, with or without chloroquine (5 μM) to block lysosomal HER2 degradation. Cells were then fixed and stained with an anti-human IgG antibody to visualize T-DM1 location (red). Phalloidin is represented in grey. **C)** Quantification of total fluorescence per cell normalized by cell area from experiment in panel B. Wilcoxon test was not significant in the WT vs KO comparison in any condition. **D)** 8-days growth curves on BT-474^WT^, BT-474^EV^ and BT-474^KO^ cells treated with DM1 25nM, T-DM1 1 μg/ml and T-DXd 1 μg/ml. Rituximab, a human IgG1 monoclonal antibody, and DMSO were used as control. **E)** Co-culture of H2B-GFP or H2B-mCherry-expressing BT-474^WT^ and BT-474^KO^ cells at baseline; stacked bar expressing population percentages over time upon treatment with T-DM1 and T-Dxd as indicated. **F)** 8-days growth curves on HCC1954^WT^, HCC1954^EV^ and HCC1954^KO^ cells treated with DM1 5nM, T-DM1 0,1 μg/ml and T-DXd 0,1 μg/ml. Rituximab, a human IgG1 monoclonal antibody, and DMSO were used as control. To draw the ribbon in log10 scale, non-positive lower boundaries (time-point 6, KO, T-DM1, time-point 8, EV, T-DM1, DM1, time-point 8, KO, T-DM1, DM1, WT, T-DM1, DM1) were substituted by the value 5000. **G)** Colony formation assay for HCC1954^WT^ and HCC1954^KO^ cell lines treated with T-DM1, DM1, T-Dxd at the indicated doses. **H)** short-term (3 days) BrdU-based dose response curves of multiple microtubule-targeting agents. Known tubulin binding sites are color-coded on the tubulin dimer structure. **I)** Immunoblot of HCC1954 cells re-expressing NF1 WT in a KO background. **J)** Active RAS pulldown of BT-474^WT^, BT-474^KO^ and BT-474^KRAS^ ^G12V^ cells.

**Supplementary figure 4.**
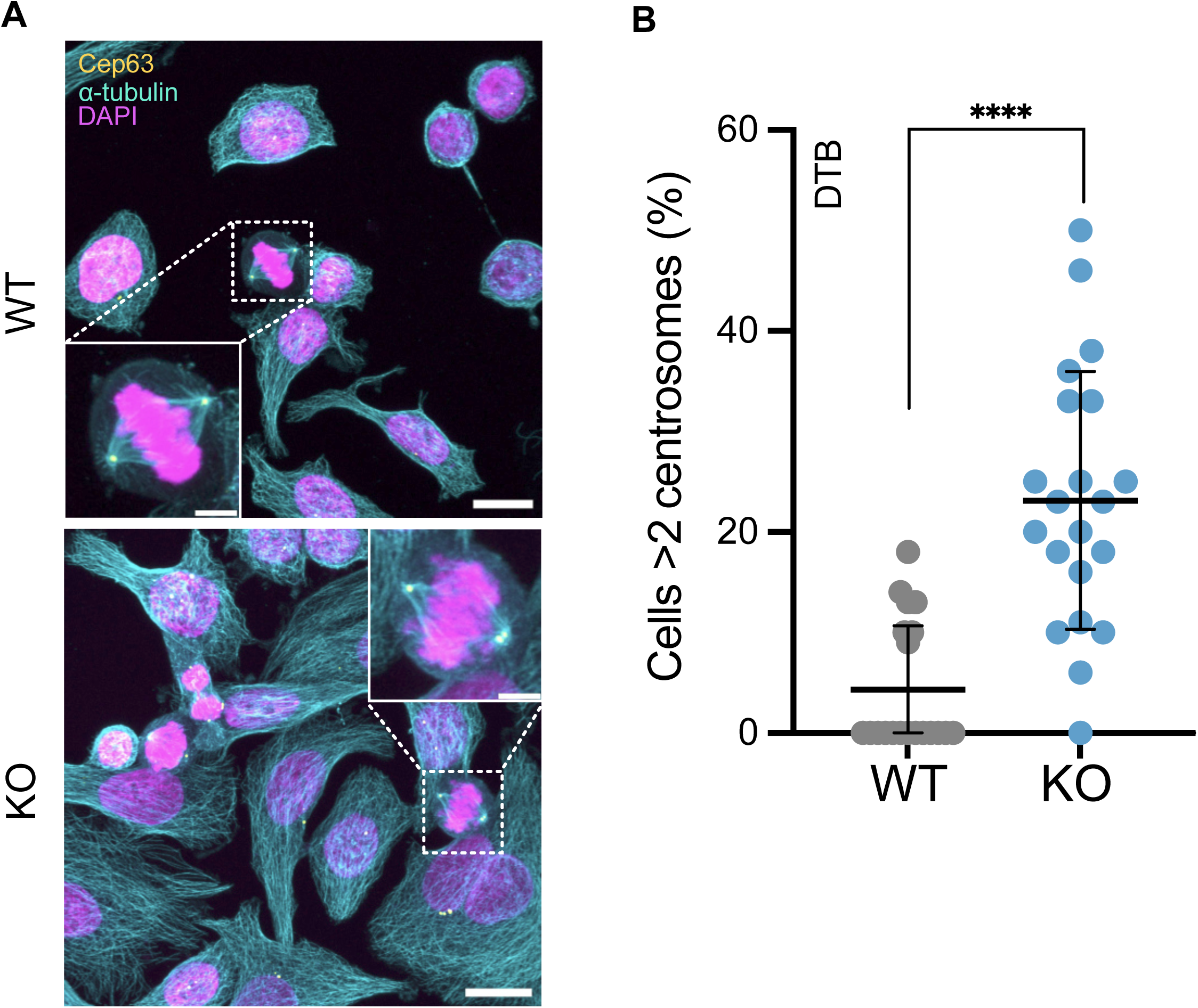
**A)** Representative images of SK-BR-3^WT^ and SK-BR-3^KO^ cells synchronized by double thymidine block. Cells are stained with anti-alpha tubulin, centrosome marker Cep63 and DAPI. Scale bar = 30 μm, scale bar = 7.5 μm. **B)** Measurement of centrosome numbers from data in panel A (mean ± SD are represented, Wilcoxon test, ****p<0.0001).

**Supplementary figure 5.**
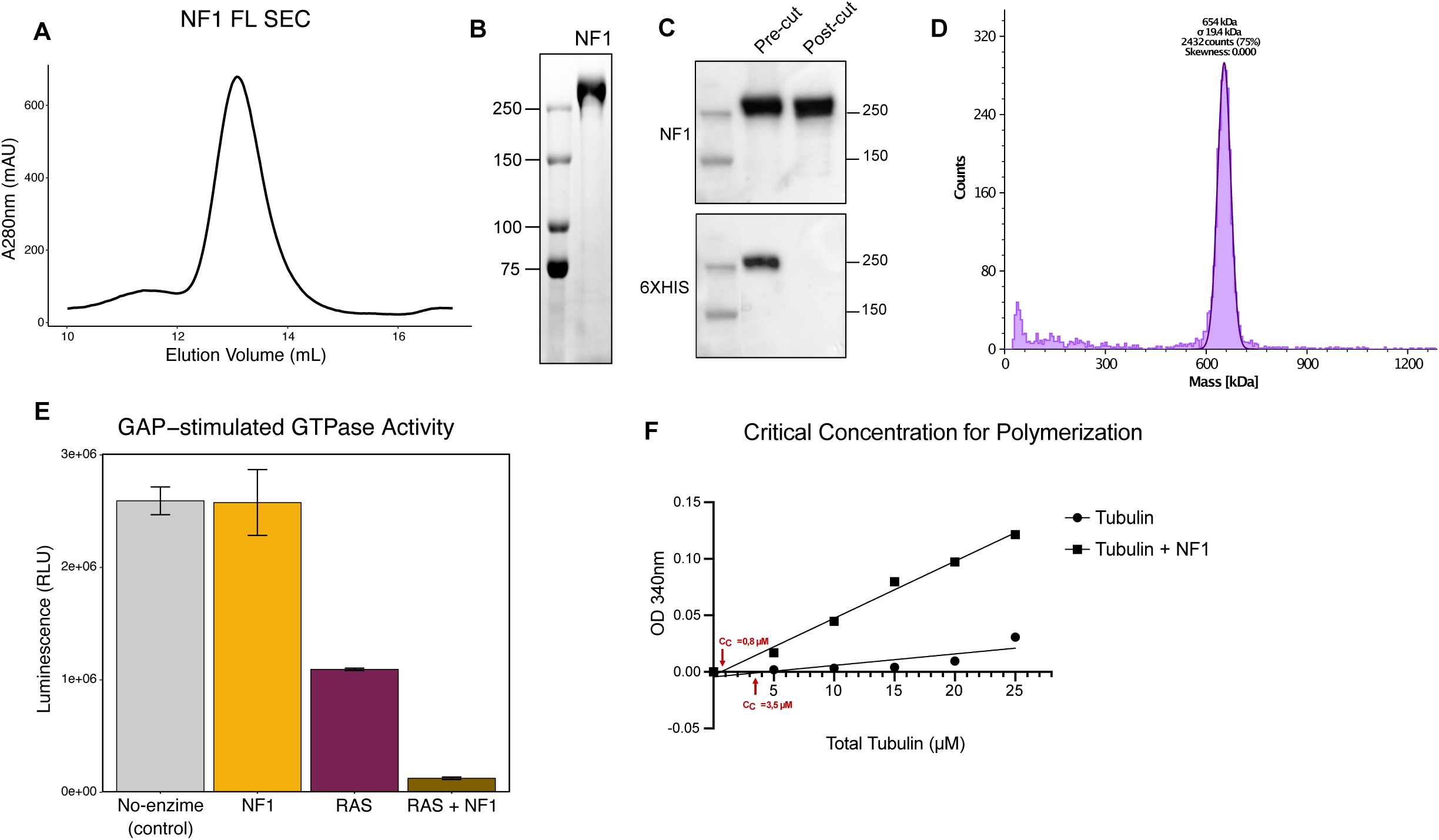
**A)** NF1 purifies as a single peak by size-exclusion chromatography. **B)** Purified recombinant NF1 is highly pure by SDS-PAGE. **C)** Western blot analysis of the purified recombinant NF1 shows the efficiency of the 6XHIS-tag cut. **D)** Mass photometry histogram. Peaks are fit by Gaussian curves. The peak is compatible with NF1 dimers. **E)** NF1 accelerates the hydrolysis of GTP by RAS. Hydrolysis was measured by GTPase-Glo kit to detect the amount of GTP remaining in the reaction after incubation. **F)** Calculation of the critical tubulin concentration for polymerization. Each point represents a reaction in which unpolymerized tubulin at the indicated concentrations was allowed to polymerize and OD recorded. We interpolated OD levels and 0 in both conditions. The intersection with X-axis is the Critical concentration (CC) (see red arrows).

**Supplementary figure 6.**
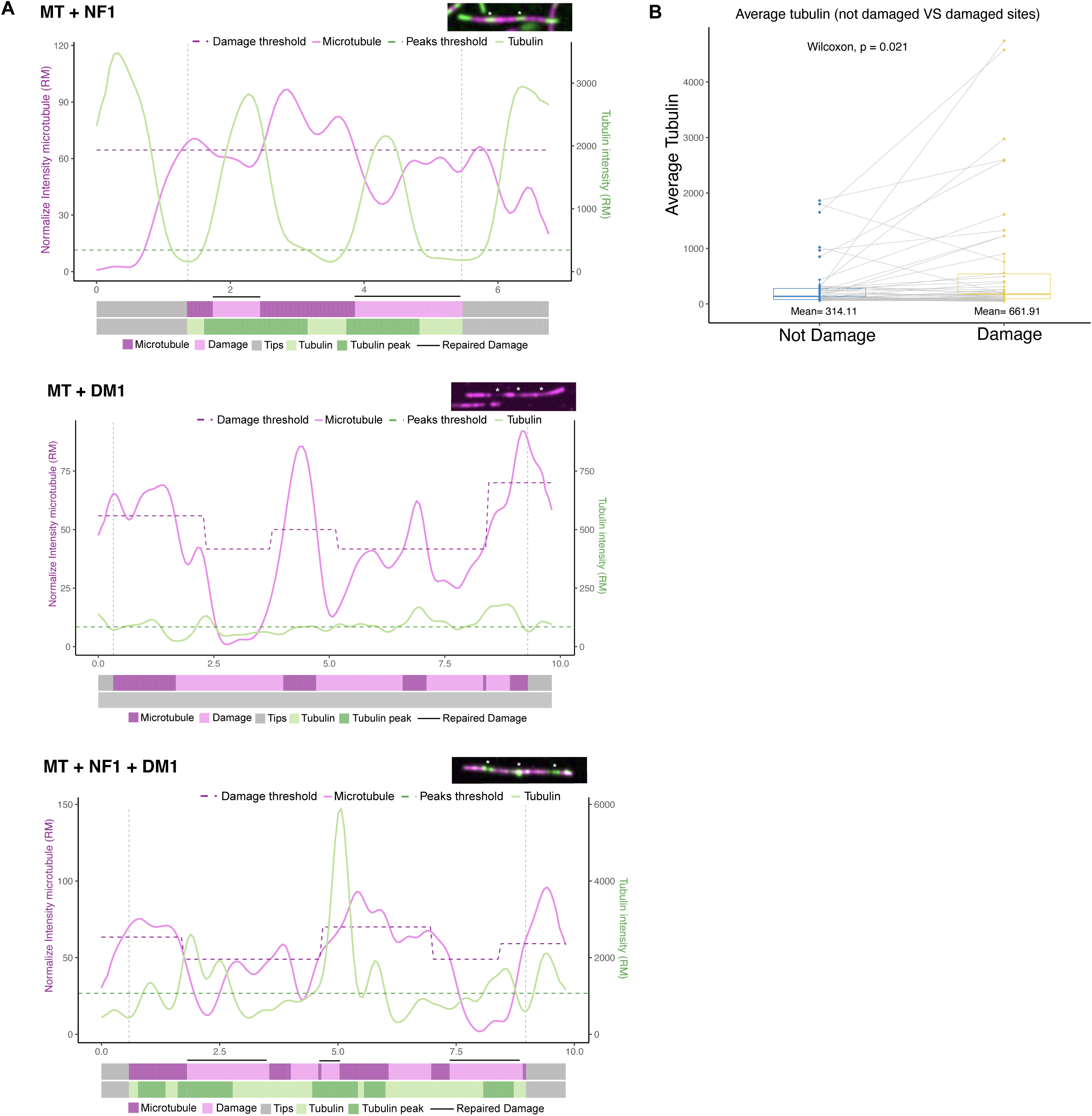
**A)** Representative plot of microtubule (magenta) and tubulin (green) signal intensity for the condition MT + NF1 300nM, MT + DM1 1μM and MT + NF1 300nM + DM1 1μM. Thresholds for the identification of damages and tubulin peaks are reported as dotted lines (violet and dark green). Horizontal bars below the intensity plot represent microtubule tips (grey), intact microtubule and damage sites (magenta); tubulin and its peaks (green). Damaged sites repaired by tubulin are highlighted with a black bar. **B)** Average tubulin intensity in non-damaged regions and in damaged sites of microtubules in the control condition. Mean values are reported, statistical significance assessed with Wilcoxon paired test *p<0.05.

**Supplementary video legends**

**Supplementary video 1**: Monitoring of cell cycle transitions in FUCCI-Ca-expressing BT-474^WT^ cells after addition of T-DM1 1 mg/ml and DRTM to monitor cell death. Cells transition through red (G1), green (S), yellow (G2/M) and white (DRTM). Cells were live-imaged for 90 hours every 10 minutes after addition of T-DM1 1 mg/ml.

**Supplementary video 2**: Monitoring of cell cycle transitions in FUCCI-Ca-expressing BT-474^KO^ cells after addition of T-DM1 1 mg/ml and DRTM to monitor cell death. Cells transition through red (G1), green (S), yellow (G2/M) and white (DRTM). Cells were live-imaged for 90 hours every 10 minutes after addition of T-DM1 1 mg/ml.

**Supplementary video 3**: Monitoring of microtubule elongation in HCC-1954^WT^ cells transiently transfected with an EB3-GFP-expressing vector and monitored by live cell imaging for 1 minute with acquisitions every 4 seconds.

**Supplementary video 4**: Monitoring of microtubule elongation in HCC-1954^KO^ cells transiently transfected with an EB3-GFP-expressing vector and monitored by live cell imaging for 1 minute with acquisitions every 4 seconds.

**Supplementary video 5**: Monitoring of microtubule elongation in HCC-1954^WT^ cells transiently transfected with an EB3-GFP-expressing vector and monitored by live cell imaging for 1 minute with acquisitions every 4 seconds. DM1 added at 0.5 nM.

**Supplementary video 6**: Monitoring of microtubule elongation in HCC-1954^KO^ cells transiently transfected with an EB3-GFP-expressing vector and monitored by live cell imaging for 1 minute with acquisitions every 4 seconds. DM1 added at 0.5 nM.

**Supplementary video 7:** Live cell imaging of BT-474^WT^ cells stably expressing H2B-GFP and monitored over 100 minutes, with acquisitions every 4 minutes. Cells were incubated with cell-permeable, fluorescent SiR-tubulin to monitor spindle formation.

**Supplementary video 8**: live cell imaging of BT-474^KO^ cells stably expressing H2B-mCherry and monitored over 100 minutes, with acquisitions every 4 minutes. Cells were incubated with cell-permeable, fluorescent SiR-tubulin to monitor spindle formation. A multipolar cell division can be appreciated.

**Supplementary video 9**: TIRF microscopy monitoring of microtubule dynamics, followed over 20 min with an acquisition every 10 seconds. GMPCPP seeds are shown in magenta, free tubulin in green. No NF1, No DM1 added.

**Supplementary video 10**: TIRF microscopy monitoring of microtubule dynamics, followed over 20 min with an acquisition every 10 seconds. GMPCPP seeds are shown in magenta, free tubulin in green. NF1 added at 100 nM.

**Supplementary video 11**: TIRF microscopy monitoring of microtubule dynamics, followed over 20 min with an acquisition every 10 seconds. GMPCPP seeds are shown in magenta, free tubulin in green. NF1 added at 300 nM.

**Supplementary video 12**: TIRF microscopy monitoring of microtubule dynamics, followed over 20 min with an acquisition every 10 seconds. GMPCPP seeds are shown in magenta, free tubulin in green. No NF1, DM1 added 1 μM.

**Supplementary video 13**: TIRF microscopy monitoring of microtubule dynamics, followed over 20 min with an acquisition every 10 seconds. GMPCPP seeds are shown in magenta, free tubulin in green. NF1 added at 100 nM, DM1 1 μM.

**Supplementary video 14**: TIRF microscopy monitoring of microtubule dynamics, followed over 20 min with an acquisition every 10 seconds. GMPCPP seeds are shown in magenta, free tubulin in green. NF1 added at 300 nM, DM1 1 μM.

## Notes

https://github.com/mazzalab-ieo/tol_NF1_scripts.

### Competing Interest Statement

The authors have declared no competing interest.

### Summary of Updates

-Title and abstract have been shortened -we have significantly expanded the last part on the in vitro assessment of microtubular damage -we have added an important retrospective analysis of patient data (figure 2L-M) -the manuscript now contains information on the ability of wild-type NF1 re-expression to rescue the KO phenotypes

## References

1. Steinmetz, M. O. & Prota, A. E. Microtubule-Targeting Agents: Strategies To Hijack the Cytoskeleton. Trends Cell Biol. 28, 776–792 (2018).

2. Tarantino, P. et al. Antibody–drug conjugates: Smart chemotherapy delivery across tumor histologies. CA. Cancer J. Clin. 72, 165–182 (2022).

3. von Arx, C. et al. The evolving therapeutic landscape of trastuzumab-drug conjugates: Future perspectives beyond HER2-positive breast cancer. Cancer Treat. Rev. 113, (2023).

4. Verma, S. et al. Trastuzumab Emtansine for HER2-Positive Advanced Breast Cancer. N. Engl. J. Med. 367, 1783–1791 (2012).

5. Cabanillas, F. et al. Phase I study of maytansine using a 3-day schedule. Cancer Treat. Rep. 62, 425– 428 (1978).

6. Prota, A. E. et al. A new tubulin-binding site and pharmacophore for microtubule-destabilizing anticancer drugs. Proc. Natl. Acad. Sci. U. S. A. 111, 13817–13821 (2014).

7. Lopus, M. et al. Maytansine and cellular metabolites of antibody-maytansinoid conjugates strongly suppress microtubule dynamics by binding to microtubules. Mol. Cancer Ther. 9, 2689–2699 (2010).

8. Oroudjev, E. et al. Maytansinoid-Antibody Conjugates Induce Mitotic Arrest by Suppressing Microtubule Dynamic Instability. Mol. Cancer Ther. 9, 2700–2713 (2010).

9. Marchuk, D. A. et al. cDNA cloning of the type 1 neurofibromatosis gene: complete sequence of the NF1 gene product. Genomics 11, 931–940 (1991).

10. Philpott, C., Tovell, H., Frayling, I. M., Cooper, D. N. & Upadhyaya, M. The NF1 somatic mutational landscape in sporadic human cancers. Hum. Genomics 11, 13 (2017).

11. Razavi, P. et al. The Genomic Landscape of Endocrine-Resistant Advanced Breast Cancers. Cancer Cell 34, 427–438.e6 (2018).

12. Smith, A. E. et al. HER2 breast cancers evade anti-HER2 therapy via a switch in driver pathway. Nat Commun 12, 6667 (2021).

13. Zheng, Z.-Y. et al. Neurofibromin Is an Estrogen Receptor-&#x3b1; Transcriptional Co-repressor in Breast Cancer. Cancer Cell 37, 387–402.e7 (2020).

14. Gutmann, D. H., Wood, D. L. & Collins, F. S. Identification of the neurofibromatosis type 1 gene product. Proc. Natl. Acad. Sci. U. S. A. 88, 9658–9662 (1991).

15. Lupton, C. J. et al. The cryo-EM structure of the human neurofibromin dimer reveals the molecular basis for neurofibromatosis type 1. Nat. Struct. Mol. Biol. 28, 982–988 (2021).

16. Naschberger, A., Baradaran, R., Rupp, B. & Carroni, M. The structure of neurofibromin isoform 2 reveals different functional states. Nature 599, 315–319 (2021).

17. Bollag, G., McCormick, F. & Clark, R. Characterization of full-length neurofibromin: tubulin inhibits Ras GAP activity. EMBO J. 12, 1923–1927 (1993).

18. Gregory, P. E. et al. Neurofibromatosis type 1 gene product (neurofibromin) associates with microtubules. Somat. Cell Mol. Genet. 19, 265–274 (1993).

19. Koliou, X., Fedonidis, C., Kalpachidou, T. & Mangoura, D. Nuclear import mechanism of neurofibromin for localization on the spindle and function in chromosome congression. J. Neurochem. 136, 78–91 (2016).

20. Bodakuntla, S., Jijumon, A. S., Villablanca, C., Gonzalez-Billault, C. & Janke, C. Microtubule-Associated Proteins: Structuring the Cytoskeleton. Trends Cell Biol. 29, 804–819 (2019).

21. Gazzola, M. et al. Microtubules self-repair in living cells. Curr. Biol. 33, 122–133.e4 (2023).

22. Triclin, S. et al. Self-repair protects microtubules from destruction by molecular motors. Nat. Mater. 20, 883–891 (2021).

23. Aher, A. et al. CLASP Mediates Microtubule Repair by Restricting Lattice Damage and Regulating Tubulin Incorporation. Curr. Biol. 30, 2175–2183.e6 (2020).

24. Lawrence, E. J., Arpag, G., Arnaiz, C. & Zanic, M. SSNA1 stabilizes dynamic microtubules and detects microtubule damage. Elife 10, e67282 (2021).

25. AACR Project GENIE Consortium. AACR Project GENIE: Powering Precision Medicine through an International Consortium. Cancer Discov. 7, 818–831 (2017).

26. Koboldt, D. C. et al. Comprehensive molecular portraits of human breast tumours. Nature 490, 61– 70 (2012).

27. Sakaue-Sawano, A. et al. Genetically Encoded Tools for Optical Dissection of the Mammalian Cell Cycle. Mol. Cell 68, 626–640.e5 (2017).

28. Stepanova, T. et al. Visualization of microtubule growth in cultured neurons via the use of EB3-GFP (end-binding protein 3-green fluorescent protein). J. Neurosci. Off. J. Soc. Neurosci. 23, 2655–2664 (2003).

29. Gulluni, F. et al. Mitotic Spindle Assembly and Genomic Stability in Breast Cancer Require PI3K-C2α Scaffolding Function. Cancer Cell 32, 444–459.e7 (2017).

30. Spurr, L. F. et al. Quantification of aneuploidy in targeted sequencing data using ASCETS. Bioinformatics 37, 2461–2463 (2021).

31. Cheng, J. et al. Accurate proteome-wide missense variant effect prediction with AlphaMissense. Science (80-.). 381, eadg7492 (2024).

32. Zhu, Z. C. et al. Interactions between EB1 and microtubules: dramatic effect of affinity tags and evidence for cooperative behavior. J. Biol. Chem. 284, 32651–32661 (2009).

33. Sherekar, M. et al. Biochemical and structural analyses reveal that the tumor suppressor neurofibromin (NF1) forms a high-affinity dimer. J. Biol. Chem. 295, 1105–1119 (2020).

34. Bollag, G. et al. Loss of NF1 results in activation of the Ras signaling pathway and leads to aberrant growth in haematopoietic cells. Nat. Genet. 12, 144–148 (1996).

35. Xu, H. & Gutmann, D. H. Mutations in the GAP-related domain impair the ability of neurofibromin to associate with microtubules. Brain Res. 759, 149–152 (1997).

36. Replogle, J. M. et al. Aneuploidy increases resistance to chemotherapeutics by antagonizing cell division. Proc. Natl. Acad. Sci. U. S. A. 117, 30566–30576 (2020).

37. Bakhoum, S. F. et al. Chromosomal instability drives metastasis through a cytosolic DNA response. Nature 553, 467–472 (2018).

38. Pearson, A. et al. Inactivating NF1 Mutations Are Enriched in Advanced Breast Cancer and Contribute to Endocrine Therapy Resistance. Clin. Cancer Res. 26, 608 LP-622 (2020).

39. Ly, N. et al. αTAT1 controls longitudinal spreading of acetylation marks from open microtubules extremities. Sci. Rep. 6, 35624 (2016).

40. Zheng, Z.-Y. et al. Neurofibromin Is an Estrogen Receptor-α Transcriptional Co-repressor in Breast Cancer. Cancer Cell 37, 387–402.e7 (2020).

41. Mosele, F. et al. Trastuzumab deruxtecan in metastatic breast cancer with variable HER2 expression: the phase 2 DAISY trial. Nat. Med. 29, 2110–2120 (2023).

42. Cheng, J. et al. Accurate proteome-wide missense variant effect prediction with AlphaMissense. Science (80-.). 381, eadg7492 (2023).

43. Favalli, V. et al. Machine learning-based reclassification of germline variants of unknown significance: The RENOVO algorithm. Am. J. Hum. Genet. 108, 682–695 (2021).

44. Tinevez, J.-Y. et al. TrackMate: An open and extensible platform for single-particle tracking. Methods 115, 80–90 (2017).

45. Ershov, D. et al. Bringing TrackMate into the era of machine-learning and deep-learning. *bioRxiv* 2021.09.03.458852 (2021).

46. Jafari, R. et al. The cellular thermal shift assay for evaluating drug target interactions in cells. Nat. Protoc. 9, 2100–2122 (2014).

47. Fink, G. et al. The mitotic kinesin-14 Ncd drives directional microtubule-microtubule sliding. Nat. Cell Biol. 11, 717–723 (2009).

