## supplementary table 1 for "NF1 modulates microtubule repair and sensitivity to antibody-drug conjugates"

**Supplementary table 1.** Summary characteristics of patients analysed in figure 2 L-M

| Variable | N | MUT, N = 5 | WT, N = 29 | p-value |
| --- | --- | --- | --- | --- |
| **HER2.status..IHC.** | 34 |  |  | 0.5 |
| 0 |  | 0 (0%) | 1 (3.4%) |  |
| 2+ |  | 0 (0%) | 5 (17%) |  |
| 3+ |  | 4 (80%) | 22 (76%) |  |
| N/A |  | 1 (20%) | 1 (3.4%) |  |
| **FISH** | 34 |  |  | 0.6 |
| Amp |  | 0 (0%) | 7 (24%) |  |
| N/A |  | 5 (100%) | 22 (76%) |  |
| **ER** | 34 |  |  | 0.5 |
| N/A |  | 1 (20%) | 1 (3.4%) |  |
| Neg |  | 3 (60%) | 19 (66%) |  |
| Pos |  | 1 (20%) | 9 (31%) |  |
| **PR** | 34 |  |  | 0.4 |
| N/A |  | 1 (20%) | 1 (3.4%) |  |
| Neg |  | 3 (60%) | 18 (62%) |  |
| Pos |  | 1 (20%) | 10 (34%) |  |
| **Prior.chemo.** | 34 |  |  | 0.3 |
| N/A |  | 1 (20%) | 1 (3.4%) |  |
| yes |  | 4 (80%) | 28 (97%) |  |
| **Prior.anti.HER2.** | 34 |  |  | 0.3 |
| N/A |  | 1 (20%) | 1 (3.4%) |  |
| yes |  | 4 (80%) | 28 (97%) |  |
| **Line.of.T.DM1** | 34 |  |  | 0.3 |
| 2 |  | 2 (40%) | 19 (66%) |  |
| 3 |  | 2 (40%) | 9 (31%) |  |
| N/A |  | 1 (20%) | 1 (3.4%) |  |
| ^1^n (%) | | | | |
| ^2^Fisher's exact test | | | | |
